# Evolution of circadian clock and light-input pathway genes in Hemiptera

**DOI:** 10.1101/2025.01.17.633578

**Authors:** Vlastimil Smykal, Hisashi Tobita, David Dolezel

**Author notes:** present address: Department of Agricultural and Environmental Biology, Graduate School of Agricultural and Life Sciences, The University of Tokyo, Bunkyo-ku, Tokyo, 113-8657, Japan (Hisashi Tobita).

## Abstract

Circadian clocks are timekeeping mechanisms that help organisms anticipate periodic alterations of day and night. These clocks are widespread, and in the case of animals, they rely on genetically related components. At the molecular level, the animal circadian clock consists of several interconnected transcription-translation feedback loops. Although the clock setup is generally conserved, some important differences exist even among various insect groups. Therefore, we decided to identify *in silico* all major clock components and closely related genes in Hemiptera. Our analyses indicate several lineage-specific alterations of the clock setup in Hemiptera, derived from gene losses observed in the complete gene set identified in the outgroup, Thysanoptera, which thus presents the insect lineage with a complete clock setup. *Nilaparvata* and Fulgoroidea, in general, lost the (6-4) photolyase, while all Hemiptera lost FBXL3, and several lineage-specific losses of dCRY and jetlag were identified. Importantly, we identified non-canonical splicing variants of *period* and *m-cry* genes, which might provide another regulatory mechanism for clock functioning. Lastly, we performed a detailed reconstruction of Hemiptera’s light input pathway genetic repertoire and explored the horizontal gene transfer of cryptochrome-DASH from plant to *Bemisia*. Altogether, this inventory reveals important trends in clock gene evolution and provides a reference for clock research in Hemiptera, including several lineages of important pest species.

**Bullet points (highlights):**

○ Evolution of clock genes (including light input pathway) was reconstructed in Hemiptera
○ New *m-cry* and *per* splicing variants were identified in certain species
○ A unique horizontal gene transfer of plant/fungal CRY-DASH was found in *Bemisia*
○ Clock setup was identified for pests: *Nilaparvata*, *Bemisia*, *Halyomorpha*, and aphids
○ Future clock research directions in Hemiptera are proposed

## 1. Introduction

Life on Earth is heavily affected by periodic changes in day and night. Consequently, nearly all organisms evolved circadian clock, an internal oscillatory mechanism measuring 24 hours (from Latin *circa…*approximately, *diem…*day), which helps them to anticipate daily changes. At a molecular level, the circadian clock of animals in general and bilateral animals in particular (i.e. mammals and insects) rely on negative transcription/translation feedback loops (TTFL) of homologous components. At least 20 clock components may be considered circadian clock proteins participating in TTFLs of animal clocks. These proteins include transcription factors, several kinases, phosphatases, proteins responsible for specific degradation pathways, etc. Although the majority of clock components are conserved across animals, certain important differences can be found.

Thanks to genetic tools and decades of effort by hundreds of scientists, the fruit fly *Drosophila melanogaster* has become the best model for the invertebrate circadian clock (Hall 2003; Hardin 2011; Ozkaya and Rosato 2012; Dolezel 2023). CYCLE (CYC) and CLOCK (CLK), two proteins from the basic helix-loop-helix (bHLH) Per-ARNT-Sim (PAS) family (Tumova et al., 2024), serve as positive elements of the feedback loop and drive mRNA expression of genes containing E-box elements in the promoter, such as *period* (*per*) and *Drosophila*-type *timeless* (*d-tim*; for the mammalian-type *tim*, *m-tim*, see text on the light input below) (Allada et al., 1998; Rutila et al., 1998). PER and dTIM proteins accumulate and enter the cell nucleus, resulting in inhibition of the transcriptional activity of the CYC-CLK dimer (Glossop et al., 1999) (Fig. 1). Although PER protein can form a homodimer (Landskron et al., 2009), the major PER-stabilizing partner is dTIM (Sehgal et al., 1994). Interaction between both proteins is key for their subcellular localization, including nuclear localization signals (NLS) and nuclear export signals (NES) (Saez et al., 1996; Singh et al., 2019; Giesecke et al., 2023). Both PER and dTIM are phosphorylated/dephosphorylated by several kinases and phosphatases contributing to their subcellular localization and stability, ultimately leading to their ubiquitination and degradation by the proteasome (Grima et al., 2002; Ko et al., 2002). Experiments on *Drosophila* PER functionally grouped phosphorylation sites into several phosphocluster regions, which based on their phosphorylation level determine PER stability and inhibitory potency and modulate temperature compensation phenotype, forming thus a phospho-switch to balance diverse PER functions throughout the day-night cycle (Kivimäe et al., 2008; Joshi et al., 2022). One of the phosphoclusters, the N-terminal phosphodegron, is responsible for PER ubiquitination and degradation in the proteasome (Chiu et al., 2008; Kivimäe et al., 2008). It includes Serine 44,45 and 47 (Chiu et al., 2008; Joshi et al., 2022), which of S47 is phosphorylated by DOUBLETIME (DBT) kinase and together with nearby phosphorylated sites, phospho-S47 generates a high-affinity atypical binding site for SUPERNUMERARY LIMBS (SLIMB) (Chiu et al., 2008), an F-box protein targeting PER for proteasomal degradation (Grima et al., 2002; Ko et al., 2002).

**Fig. 1.**
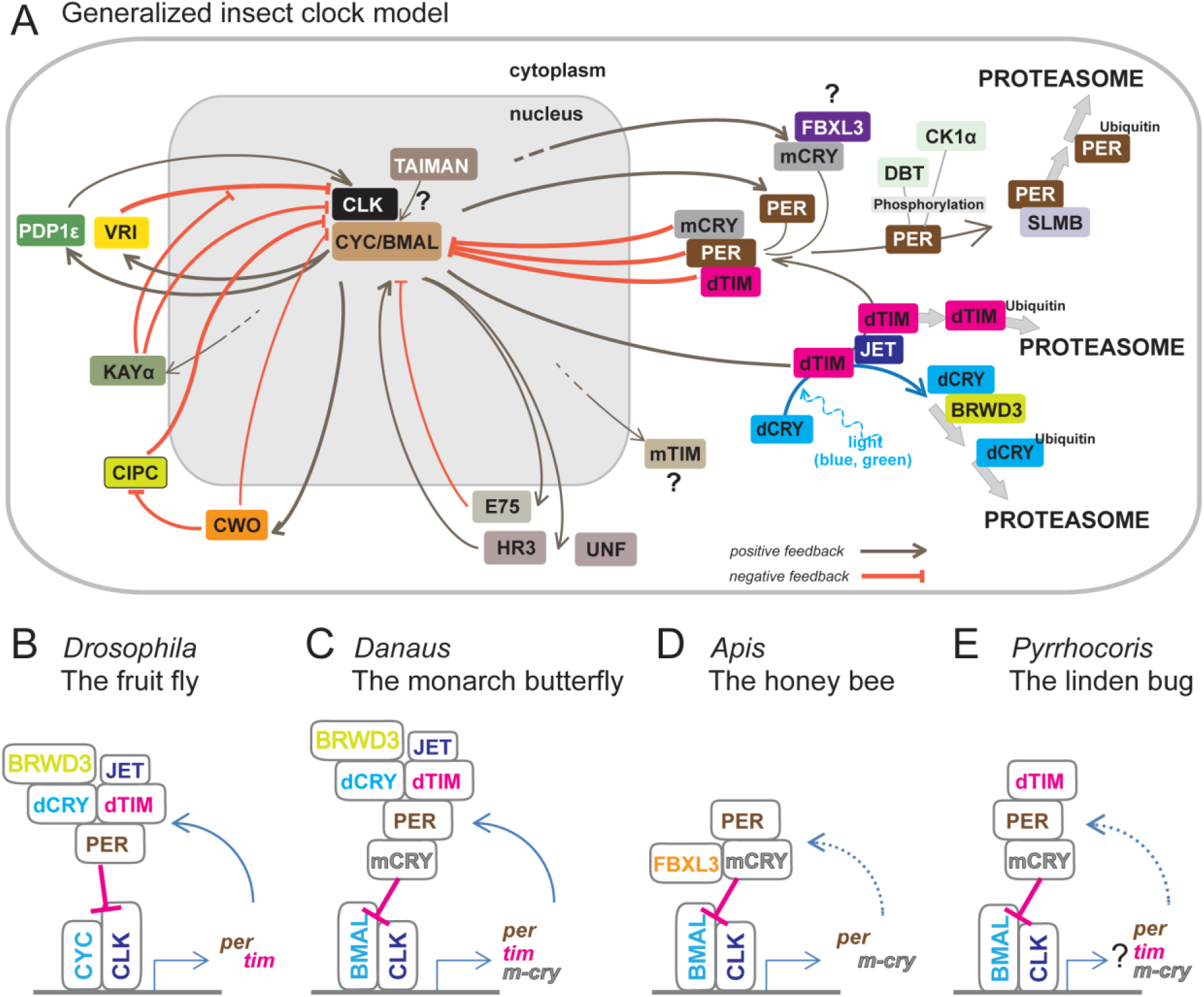
(A) Simplified summary of the most variable components in insect clockwork models. The generally conserved positive components, CLOCK (CLK) and BMAL/CYCLE (CYC) drive the expression of negative feedback loop components. Adapted from Doležel 2023. **(B)** In *Drosophila*, key negative components include PER and dTIM. The light-induced degradation of these components requires dCRY, JET, and BRWD3. **(C)** In addition to *Drosophila* components, the monarch butterfly possesses mammalian-type CRY (mCRY), which acts as the primary repressor. **(D)** The honeybee has lost dTIM, dCRY, and JET. **(E)** The clock in the linden bug relies predominantly on mCRY and PER, with some contribution from dTIM.

As the most thoroughly studied clock genes, *per* and *d-tim*, can illustrate the fine-tuning role of alternative splicing in circadian clock regulation. At low temperatures, efficient splicing of an intron located in the 3’ untranslated region (UTR) of *per* results in advance accumulation of PER protein, which further manifests in advanced evening activity typical for fruit flies at low temperatures (Majercak et al., 1999, 2004). A complex splicing pattern has been reported for *d-tim* alternative splicing. For example, alternative retention of an intron located in the center of the gene at high temperatures results in mRNA that is likely destroyed by a non-sense mediated decay mechanism (NMD) and can only encode unstable and nonfunctional protein (Martin Anduaga et al., 2019; Foley et al., 2019; Shakhmantsir et al., 2018). This splicing might be one of many adjustments cumulatively resulting in a temperature-compensated clock in *Drosophila*.

The circadian oscillation is further regulated by the action of Clock-Interacting Protein Circadian (CIPC) and CLOCKWORK ORANGE (CWO). First, when CLK-CYC is released from E-boxes, CWO binds to E-boxes and reinforces the repression (Zhou et al., 2016). At the same time, CWO simultaneously represses the expression of *Cipc*, which itself is a repressor of CLK-CYC transcription (Rivas et al., 2021). Thus, while CWO represses CLK-CYC, it also activates their transcription later by repressing their repressor, CIPC, altogether explaining the seemingly contradicting roles of simultaneous activator and repressor (Kadener et al., 2007; Richier et al., 2008).

Another group of TTFL involves basic leucine zipper (bZIP) proteins VRILLE (VRI), PAR DOMAIN PROTEIN 1 (PDP1), and KAYAK (KAY). Their genes, specifically certain isoforms, contain E-boxes that are recognized by the CLK-CYC dimer and drive the periodic mRNA transcription. First, the mRNA of *vrille* (*vri*) peaks, whereas the *Pdp1-epsilon* isoform reaches its maximum several hours later, and a similar delay is observed in protein abundance. VRI serves as a transcription repressor that recognizes D-boxes (also known as V/P boxes), cis-regulatory sequences located in the *Clk* promoter (Blau and Young, 1999; Cyran et al., 2003). The role of VRI is gradually mitigated by two additional bZIP proteins. KAY, a homolog of mammalian FOS, directly binds VRI and inhibits its repressive role (Ling et al., 2012); whereas PDP1 recognizes and binds to the same DNA sequence. However, PDP1 serves as a transcriptional activator; thus, the actions of the three bZIP proteins described here result in periodic expression of *Clk* mRNA that is in antiphase to *per* and *d-tim* oscillation in *Drosophila* (Cyran et al., 2003).

The mammalian clock relies on a TTFL with RORα and REV-ERBα, two orphan nuclear receptors cyclically expressed by CLK-BMAL (Brain and Muscle ARNT-like 1, a mammalian homolog of CYC). Similarly, *Drosophila* clock machinery involves nuclear hormone receptors E75 (homolog of REV-ERBα) and UNFULFILLED (UNF) (Jaumouille et al., 2015). Interestingly, the role of UNF seems to be a *Drosophila*-specific clock idiosyncrasy, as this protein has not been identified as a clock component in other insect models. Instead, HR3 (homolog of RORα) was identified by RNA interference in an early-branching insect species, the firebrat *Thermobia domestica* (Kamae et al., 2014).

TAIMAN (TAI), a bHLH-PAS protein also known as steroid receptor coactivator (SRC) or Nuclear Receptor Coactivator (NCoA), was recognized as a new clock component in two distant insect lineages: true bugs and cockroaches (Smykal et al., 2023). TAI, initially identified as a developmental gene in a forward genetic screen (Bai et al., 2000), participates in numerous developmental processes, including metamorphosis, reproduction, and adult diapause (Smykal et al., 2014a; Smykal et al., 2014b; Hejnikova et al., 2022). Its roles involve participation in juvenile hormone and ecdysteroid receptions, but its function in the circadian clock seems to be independent of either of these hormones. Importantly, the mammalian ortholog SRC2 participates in mouse circadian rhythmicity, while SRC3 is involved in ultradian metabolic rhythms, which further underscores the evolutionarily conserved participation of TAI in the clock (Stashi et al., 2014; Meng et al., 2022). The role of TAI in the *Drosophila* circadian clock remains elusive.

As already illustrated regarding nuclear receptors (Kamae et al., 2014; Jaumouille et al., 2015), there are specific modifications in the clock setup related to particular insect lineages. Additional major lineage-specific changes involve the dTIM protein and its partners. Similar to the mammalian clock, dTIM has been lost in Hymenoptera (Rubin et al., 2006) and independently in termites (Kotwica-Rolinska et al., 2022a). Notably, this gene loss is further accompanied by the loss of *Drosophila*-type *cryptochrome*, *d-cry* (in the *Drosophila* literature often annotated as *cry*, in some evolutionary comparisons annotated as *cry1* or *cryI*), a gene encoding protein related to DNA photolyases that lacks the DNA-repair activity (Yuan et al., 2007; DeOliveira and Crane, 2024). Another homolog of DNA photolyases, the mammalian-type *cry*, *m-cry* (annotated as *cry2* or *cryII* in some evolutionary comparisons), has been lost in *Drosophila*. The mCRY protein, first identified as a major component of the clock in vertebrates (Kume et al., 1999; Putker et al., 2021), is a key component of clock machinery in numerous insects, including the monarch butterfly, cockroaches, and true bugs (Ikeno 2011a; Bazalova et al., 2016; Zhang et al., 2017; Werckenthin et al., 2020; Kotwica-Rolinska et al., 2022a).

Although dCRY contributes to the clock oscillation and robustness in Lepidoptera and to some extent even in *Drosophila* (Dolezelova et al., 2007; Iiams et al., 2024; Tobita and Kiuchi, 2024), particularly in the peripheral oscillator (Collins et al., 2006), its major role in the fruit fly is in the light-mediated entrainment, where dCRY serves as a deep brain photoreceptor (Emery et al., 2000). Upon blue light illumination, dCRY directly interacts with dTIM, and both proteins are targeted for proteasomal degradation when JETLAG (JET) and BRWD3/Ramshackle play key roles (Ceriani et al., 1999; Koh et al., 2006; Peschel et al., 2009; Ozturk et al., 2013). While the latter is a developmental gene (D’Costa et al., 2006), JET has been lost in several insect lineages (Kotwica-Rolinska et al., 2022a), and often, but not always, the loss accompanies either the loss of dCRY or major modification of dTIM (Bullo et al., 2024). Interestingly, dCRY-independent light input affecting dTIM has been reported for the QUASIMODO (QSM), a membrane-anchored *Zona pellucida* domain protein (Chen et al., 2011). The detailed mechanism of QSM action in the *Drosophila* clock remains elusive and so is the role of this protein in the clock of other species.

In addition to dCRY-dTIM entrainment, light information accesses the *Drosophila* clock via canonical visual opsins (Saint-Charles et al., 2016). These photoreceptors are found in the compound eyes, ocelli, and the Hofbauer-Buchner eyelet, which is located underneath the compound eye and projects to the brain pacemaker (Helfrich-Förster et al., 2001; Helfrich-Förster 2019). Their phototransduction cascade includes two phospholipases C-β encoded by the *no receptor potential A* (*norpA*) gene and PLC-β encoded by the *Plc21C* gene (Helfrich-Forster et al., 2001; Ogueta et al., 2018). In crickets, compound eyes are the only circadian photoreceptors. Both the opsins, particularly the green-sensitive one, and cooperation between *cryptochromes* and *c-fos* mediate photic entrainment of the circadian clock (Komada et al., 2015; Kutaragi et al., 2018). Similarly, the retinal cells in the compound eyes are key for the light input into the photoperiodic timer in the bean bug *Riptortus pedestris* (Xi et al., 2017). This contrasts with the localization of the light-sensitive regions to the brain in aphids, as was elegantly shown using a capillary-focused light source (Lees 1964). However, the actual photoreceptor is not clearly pointed because aphids possess dCRY, dTIM, and opsins (Cortés et al., 2010; Collantes-Alegre et al., 2017). Thus, even more closely related groups (such as aphids and bean bugs) may differ in the anatomy of their light input.

As briefly illustrated above and in literature (Tomioka 2014; Tomioka and Matsumoto 2015; Tomioka and Matsumoto 2019; Kotwica-Rolinska et al., 2022a; Dolezel 2023), certain important differences exist in the clock setup among various insect lineages. Hemiptera is a large insect order and together with Psocodea and Thysanoptera form Hemipteroid insects (Paraneoptera) (Johnson et al., 2018). This large assemblage contributes to more than 10% of insects’ diversity and with numerous pests and disease vectors they are important both from basic research and practical perspectives. The circadian clock of Hemiptera was explored to a different extent in some species. In *Rhodnius prolixus*, the anatomy of the clock was studied in the neuroanatomical context and from the developmental perspective (Vafopoulou et al., 2007; Vafopoulou and Steel, 2012). Similarly, the localization of circadian clock products was assessed using *in situ* hybridization (*m-cry*, *d-cry*, *per* and *d-tim* transcripts) and immunohistochemistry (dCRY and PER antibodies) in the pea aphid *A. pisum,* and analyzed in the context of possible seasonality-relevant outputs, such as insulin-like proteins (Colizzi et al., 2021; Barberà et al., 2017, 2022). Furthermore, splicing isoforms were analyzed with temporal resolution under both short and long photoperiods, however, no association of specific transcript variants to a particular aphid strain or photoperiod could be identified (Barberà et al., 2017). At a functional level, the circadian clock genes were analyzed in two hemipteran model species, the bean bug *R. pedestris* and the linden bug *P. apterus*. In both species, systemic RNA-mediated interference (RNAi) is routinely used. Furthermore, in the linden bug, CRISPR-Cas9 gene editing was established and used to fully explore the role of several clock genes (Kotwica-Rolinska et al., 2019, 2022a, 2022b).

Given the above-described complexity of circadian clocks in insects, the inventory of clock genes in Hemiptera promises to identify important changes in the clock setup of several agricultural pests including aphids, psyllids, true bugs, and brown plant hoppers. Revealing lineage-specific traits unique to this diverse group of pests, or identifying more general patterns within Hemiptera, is essential, as these traits may influence their behavior, including the daily activity cycles, regulation of seasonality, and reproduction. Additionally, these genes are likely involved in migratory behaviors, which can significantly affect how many pest species move between habitats, especially in response to seasonal variations. Therefore, a comprehensive inventory of circadian clock genes is necessary to elucidate these functional relationships, paving the way for more effective pest management strategies that consider the intricate biological and ecological interplay dictated by circadian mechanisms.

## 2. Methods and materials

### 2.1 Dataset and gene discovery

We used a similar approach to identify genes in selected insect lineages as in Smykal et al. (2020). In brief, the initial steps of gene identification involved a BLASTP search in the protein database, a TBLASTN search in transcriptome shotgun assemblies (TSA), and we also used keywords if the genome/protein dataset was annotated. Importantly, several hits from the same species (usually more than five) were retrieved for the initial analysis. All retrieved sequences were first aligned using MAFFT, and fast tree analysis was used to identify duplicates, redundancies, and non-clock sequences. Sequences were further verified by reciprocal BLAST (when the retrieved sequences served as a query in new searches) and/or by manual inspection of alignments. Several rounds of refined and reciprocal searches were performed in particular lineages and species to retrieve all available sequences.

### 2.2 Phylogenetic analyses

Sequences were aligned using the MAFFT E-ins-i algorithm and phylogenetic trees were inferred using RAxML algorithm (Geneious Prime). For bHLH-PAS proteins, we used as reference *Apis*, *Thrips*, *Drosophila*, *Danaus*, and *Halyomorpha* sequences from the recently published dataset (Tumova et al., 2024). To reconstruct the evolution of FBXL proteins, we utilized sequences from our early studies and a recent analysis of the cricket FBXL repertoire (Takeuchi et al., 2023). As a reference of casein kinases, we used sequences from (Thakkar et al., 2022), and for the remaining clock genes, we used the linden bug *P. apterus* (Kotwica-Rolinska et al., 2022a) and *Drosophila*.

### 2.3 Gene loss

Whenever we failed to identify a clock gene in a particular species, we attempted to clarify whether this absence reflects insufficient sequencing (a very common situation especially in TSA), poor genome annotation, or if a gene loss is a likely explanation. As gene loss, we consider the situation when the gene is absent in all species of a monophyletic group for which well-sequenced genomes and transcriptomes are available2.4 Gene synteny

To reconstruct gene synteny, we used the same approach as previously (Smykal and Dolezel, 2023). First, we prospected genomic (*6-4) photolyase* locus in *Homalodisca vitripennis* (Auchenorrhyncha, Membracoidea superfamily) and genomic *cry-DASH* locus in *Bemisia tabaci* to identify protein-coding upstream and downstream syntenic genes. The syntenic genes’ coding sequences were then searched in TSA (Transcriptome Shotgun Assembly) databases of *Nilaparvata lugens* and *Laodelphax striatellus* (both Auchenorrhyncha, Fulgoroidea superfamily), or *Trialeurodes vaporariorum* (Sternorrhyncha, Aleyrodidae family), respectively, by BLAST (blastn, tblasn). The TSAs were then used as a query to identify genomic contigs/scaffolds with syntenic genes and to localize syntenic genes’ positions and the overall organization within the genomic regions. The number of protein-coding genes between syntenic genes for *N. lugens* and *L. striatellus* contigs is based on the original genome annotation. The *cry-DASH*-harboring region in Fig. 5 was BLASTED against GenBank to identify the source of the closest horizontal gene transfer donor. Syntenic gene IDs for Fig. 5 (*phr6-4*) and Fig. 6 (*cry-DASH*) are in Supplementary Table S1 and Table S2, respectively. The resulting syntenies were drawn in CorelDRAW 6X (Alludo, Ottawa, Canada).

### 2.5 Gene isoforms and gene models

To identify alternatively spliced isoforms, we used a similar approach as in (Smykal et al., 2023), when we explored *Pyrrhocoris apterus* whole mRNA Oxford Nanopore Technology (ONT) reads. We used the longest *m-cry* and *per* coding sequences (*period*: MW662133.1; *m-cry*: MW662132.1), blasted them against our custom-made *P. apterus* ONT brain transcriptomic databases and retrieved all mRNA reads. The reads were mapped to the in-house *P. apterus* genome using Minimap2 within Geneious Prime, allowing us to reconstruct the gene structure, determine the splice isoforms, and estimate their expression ratio. *N. lugens* and *H. halys per* and *m-cry* isoforms were primarily obtained from the annotated genomes, with some support from species-specific TSA databases. Protein domains (DNA_photolyase, FAD_binding_7, PAS, PAS_11, PeriodC) were predicted by using the PFAM database (Mistry et al., 2021). The Nuclear Localization Sequences (NLS) were predicted using NLStradamus or Eukaryotic Linear Motif (ELM) resources (Nguyen Ba et al., 2009; Kumar et al., 2022). The gene isoforms and gene model figures were drawn in CorelDRAW 6X.

## 3. Results

### 3.1. The ancestral clock

First, we attempted to identify homologs of clock components known in insects (Dolezel 2023; Tomioka and Matsumoto, 2015). Although this study focused on the clock gene inventory in the brown planthopper *Nilaparvata lugens*, we decided to interpret this inventory in the broader context of the entire group Hemiptera. As an outgroup, two thrips (Thysanoptera) species (*Frankliniella occidentalis* and *Thrips palmi*) and a body louse (*Pediculus humanus*) were used. As additional reference insects, we included the fruit fly *Drosophila melanogaster*, the black soldier fly *Hermetia illucens*, the red flour beetle *Tribolium castaneum*, the monarch butterfly *Danaus plexippus*, and the honeybee *Apis mellifera* (Fig. 2-4).

**Fig. 2.**
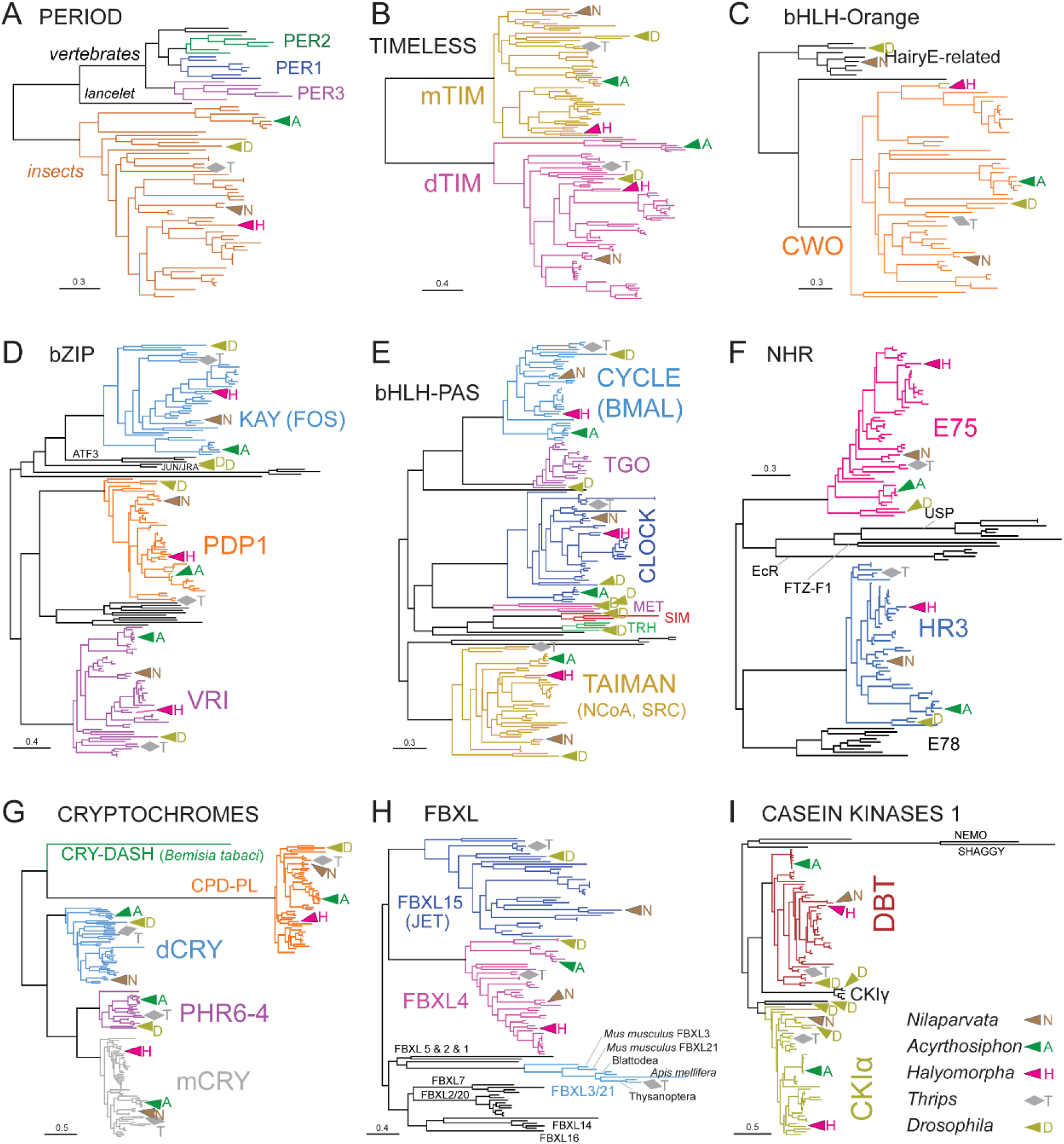
Phylogeny of circadian clock proteins. (A) A phylogenetic tree illustrates the relatedness of insect PER to vertebrate PER1, PER2, and PER3. (B) *Drosophila*-type timeless (dTIM) and mammalian-type timeless (mTIM) are clearly distinguishable. (C) bHLH-Orange transcription factors. (D) Transcription factors with the basic leucine zipper domain (bZIP) include three clock components: Vrille (VRI), Par domain protein 1 (Pdp1), and Kayak (Kay)/Fos, which are well-separated from each other and related to ATF3 and JUN proteins. (E) Proteins with the basic helix-loop-helix PER-ARNT-SIM (bHLH-PAS) domain include three circadian clock components: CYCLE/BMAL, CLOCK, and TAIMAN (also known as NCoA or SRC). (F) Nuclear Hormone Receptors (NHR) include two circadian clock components: HR3 and E75. (G) Insect photolyases and cryptochromes form five clusters; however, the *cry-DASH* gene is found only in *Bemisia*. (H) Although the phylogeny of all F-box and leucine-rich repeat proteins (FBXL) is not easily reconstructed, the three proteins relevant to the circadian clock are well-defined. (I) Doubletime (DBT), Casein Kinase Iα (CKIα), and CKIγ form well-separated clades. Note duplications CKIα-like in *Drosophila*. NEMO and SHAGGY proteins were used as an outgroup.

Consistent with early studies (Yuan et al., 2007), the ancestral combination of clock components includes PER, dTIM, mCRY, and dCRY. This clock setup, characteristic to Lepidoptera, was identified in Thysanoptera, in which also FBXL3 protein was identified (Kotwica-Rolinska et al., 2022a) (Fig. 1). FBXL3 and its close paralog FBXL21 participate in mammalian clock where these proteins bind to mCRYs (Godinho et al., 2007; Siepka et al., 2007; Hirano et al., 2013). Thus, Thysanoptera seems to represent one of a few insect lineages with a complete set of circadian genes. However, the actual functional role of these genes has not been addressed.

### 3.2. Various reduction of clock components in outgroup species

The additional, more distant outgroups than Thysanoptera, characterize the diversity of clock setups identified in insects. The body louse *Pediculus humanus* is an example of a species in which dCRY and JET have been lost. A similar loss was reported for *A. mellifera* and *T. castaneum*, however, more detailed species sampling revealed that all three losses of dCRY and JET are independent (Kotwica-Rolinska et al., 2022a). Furthermore, the honeybee (and all Hymenoptera) also lost dTIM and its clock setup, thus, resembling the mammalian one (Rubin et al., 2006). The lepidopteran clock represented by the monarch butterfly *Danaus plexippus* contains the nearly complete clock gene toolkit when only FBXL3 is missing.

The fruit fly *Drosophila* has lost mCRY and contains dCRY (in addition to (6-4) photolyase and CPD photolyase). Interestingly, *Hermetia illucens*, a dipteran species belonging to (infraorder) Stratiomyomorpha contains both genes. Thus, the loss of mCRY seems to be confined to a relatively narrow group of Cyclorrhapha (Muscomorpha).

### 3.3. Evolution of clock setup in Hemiptera

Hemiptera is an assemblage of three groups, usually recognized as orders: Auchenorrhyncha, Sternorrhyncha, and Heteroptera. The phylogenetic relationships among them appear to be well resolved, thanks to modern phylogenomic approaches. The separation of Hemiptera from Thysanoptera is estimated to have occurred approximately 400 million years ago (Johnson et al., 2018). As illustrated in Fig. 4, the majority of circadian clock genes are present in all Hemiptera. An exception is *d-cry*, which has been lost in *Halyomorpha* (Pentatomomorpha) and *Cimex* (Cimicomorpha). This loss is accompanied by the loss of *jetlag* (*fbxl15*). An independent loss of *jetlag* is observed in Aphidoidea (aphids and phylloxera species); however, these lineages possess *d-tim*. To clearly identify the *m-cry* gene, we have also annotated its related paralog, (*6-4) photolyase* (*phr6-4*). This gene, which does not play a role in the circadian clock, has been lost independently in true bugs (Heteroptera) and Fulgoroidea (Fig. 5).

### 3.4. Evolution of opsin genes and norpA

Phylogenetic reconstruction of insect opsins revealed five well-separated groups (Fig. 3A). For simplicity, we have adopted the nomenclature used in a previous aphid-focused study (Collantes-Alegre et al., 2018) to label these groups. The branches corresponding to Long-Wavelength Opsins (LWO), Rhodopsin 7 (Rh-7), Short-Wavelength Opsins (SWO), and Medium-Wavelength Opsins (MWO) contain only insect sequences.

**Fig. 3.**
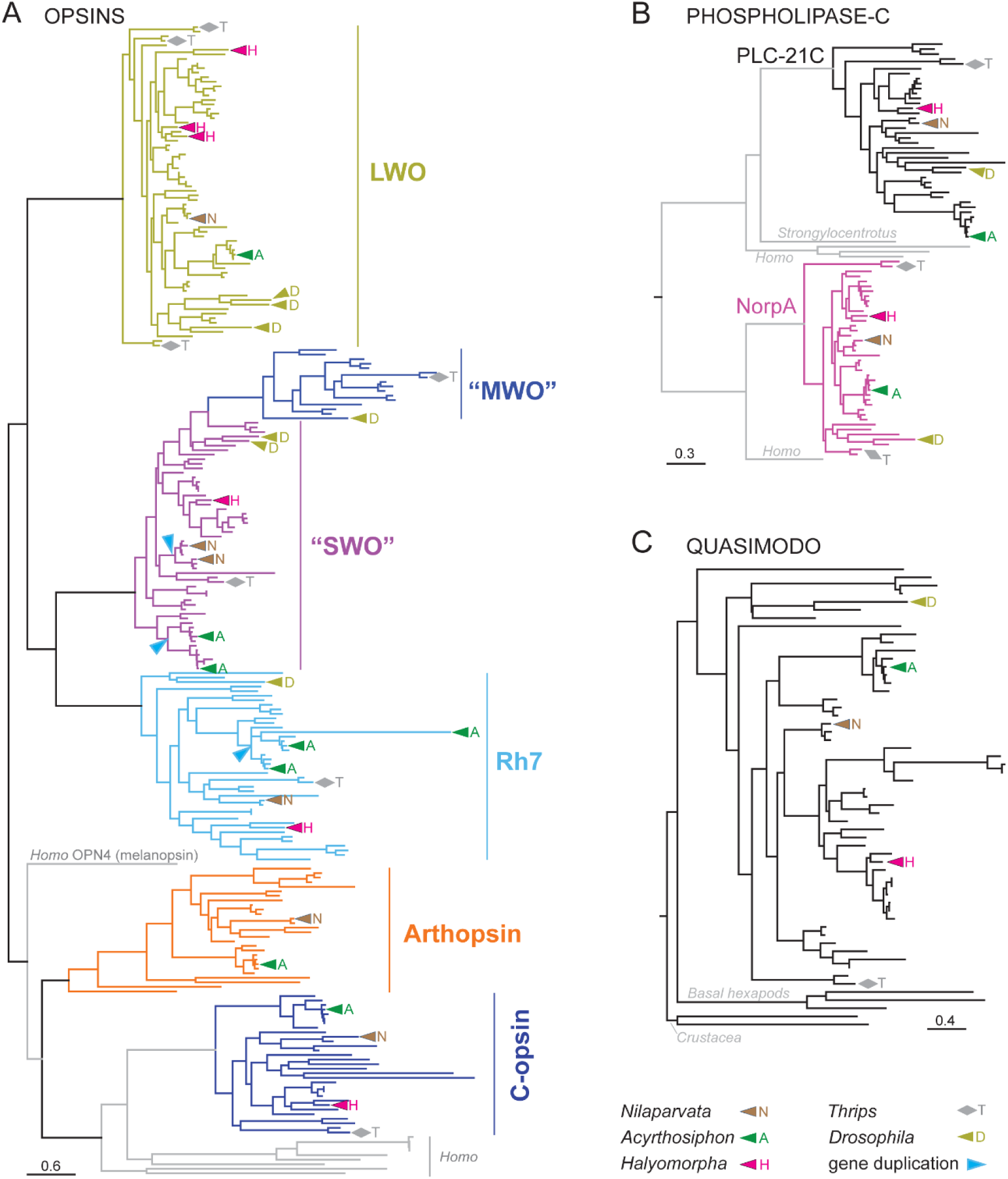
(A) Insect opsins form five well-separated monophyletic groups described as follows: Long wavelength opsin (LWO); a monophyletic group comprising medium (MWO) and short (SWO) wavelength opsins; the rhodopsin 7 (Rh7) group; Arthropsin; and C-opsin. However, the group names reflect the names of proteins in representative species, whereas the actual wavelength sensitivities remain mostly unknown. A turquoise arrowhead highlights the SWO gene duplication observed in planthoppers (upper arrowhead) and aphids (lower arrowhead). (B) Phylogeny of Phospholipase C-21 (PLC-21C) and Phospholipase C-β (PLC-β), the latter known as No Receptor Potential A (NorpA). (C) Insect Quasimodo (QSM) phylogeny, where QSM from Basal hexapods and Crustacea (*Homarus* and *Daphnia*) served as outgroups.

The distinction between the SWO and MWO groups in the phylogenetic tree is somewhat unclear and arbitrary (see Discussion for further details). To emphasize this ambiguity, we use quotation marks for these groups (“SWO” and “MWO”). Notably, three heteropteran groups have lost the “MWO” opsin (Fig. 4). Interestingly, independent duplications of “SWO” have occurred in two of these groups, Fulgoroidea and Aphidoidea. Collantes-Alegre et al. (2018) suggested that an amino acid substitution from K90 to V90 in transmembrane domain 2 (TM2) could have caused a wavelength shift in one SWO in aphids. Such neofunctionalization might explain aphid sensitivity to medium wavelengths, even though no aphid opsin branches are found within the MWO cluster. Similarly, an independent duplication of the SWO gene in Fulgoroidea coincides with the loss of the MWO group. Furthermore, these SWO paralogs in Fulgoroidea are distinguished by a T-to-I substitution, located one amino acid downstream from the K90-to-V90 change observed in aphids (Supplementary Figure S1).

Rh-7 opsins are present in all hemipteran species and in most reference insect species. They have been lost only in the honeybee and the red flour beetle. Among the five analyzed aphid species, four contain two Rh-7 genes. Closer examination suggests that this is the result of a specific duplication event (Fig. 3A). Additionally, the paralogous proteins differ at several amino acid positions (Supplementary Fig. S1).

Human OPN4, which encodes melanopsin, branches between the previously described insect opsin groups (LWO, SWO, MWO, and Rh-7) and two additional groups: Arthropsins and C-opsins. Arthropsins have been lost in Thysanoptera, Psocodea, and holometabolous outgroups (Fig. 4). Within Hemiptera, Arthropsins have only been lost in *Halyomorpha* (Pentatomomorpha). Ciliate opsins (C-opsins) branch as a sister group to Human OPN1 proteins (five paralogs), RHO, and OPN3. Two additional human protein sequences, OPN5 and peropsin, branch between C-opsins and Arthropsins. The final human protein, OPN4, which encodes melanopsin, branches between the Arthropsin/C-opsin group and the remaining insect opsins (LWO, MWO, SWO, and Rh-7; Fig. 3A).

In addition, several single-species gene duplications were identified, such as two Arthropsin genes in *Homalodisca* and two Rhodopsin 7 genes in *Apolygus*. However, these cases were not investigated further, and no attempt was made to determine whether they represent true gene duplications or errors in genome assembly. Two phospholipases, PLC-21C and NorpA, are relevant for circadian clock in *Drosophila*. Interestingly, the *norpA* gene has been duplicated in Thysanoptera, which is, to our knowledge, the only documented instance of *norpA* gene duplication in insects (Fig. 3B).

3.5. Unique cry-DASH in *Bemisia*

CRY-DASH-type cryptochrome is absent in all insects except the silverleaf whitefly *Bemisia tabaci*. The phylogenetic analysis placed *B. tabaci* CRY-DASH, encoded by an intron-less gene, among plant and fungi CRY-DASH proteins and suggested a horizontal gene transfer (HGT) as a possible origin (Kotwica-Rolinska et al., 2022a). We inspected the *B. tabaci cry-DASH* locus and its flanking regions for a trace of donor, non-insect DNA. We first searched for seven protein-coding genes upstream and downstream of *B. tabaci cry-DASH* in the genome of related (but *cry-DASH*-less) whitefly *Trialeurodes vaporariorum* to estimate the size of the genomic region transferred from a host. Although spread over >47 Mbp, all seven genes were successfully identified and localized in the *T. vaporariorum* contig VMOF01000024.1. Roughly ∼ 130 kbp-long *cry-DASH* genomic region between syntenic genes was blasted against GenBank genome databases. The 33-kbp-long *Solenum lycopersicum* genomic contig was the only long positive hit, mapping ∼ 21 kbp downstream to the *cry-DASH* gene (Fig. 6A). The corresponding *B. tabaci* genomic sequence is ∼90% identical to *S. lycopersicum* contig with two suspicious short protein-coding genes unique to *B. tabaci* and a peculiar CANIN domain-containing gene (Fig. 6B). The *S. lycopersicum* genomic fragment is not mapped as a single uninterrupted fragment but as seven ≥700-bp-long fragments, suggesting further evolution of the locus in *B. tabaci*. Importantly, *S. lycopersicum* DASH is not localized within the 33-kbp DNA contig. Additionally, a comparison of coding sequences revealed only ∼40% identity between *B. tabaci* and *S. lycopersicum* CRY-DASH, in contrast to the ∼90% identity observed in downstream-located DNA. Previous analyses suggested that *Bemisia* CRY-DASH is related to plants or fungi; however, the phylogenetic relationship had not been reconstructed in detail. These discrepancies prompted us to expand the CRY-DASH sequence dataset and reanalyze the phylogenetic relationships (Fig. 6C). *B. tabaci* CRY-DASH clusters with fungal CRY-DASH proteins from several fungal lineages, particularly Ascomycota, and is clearly separated from the strictly plant branch. The presence of four plant CRY-DASH sequences identified in the Transcriptome Shotgun Assembly (TSA) databases within the ‘fungal’ cluster is highly suspicious and may represent cross-contamination of plant samples with fungal material (Fig. 6C). Although the cry-DASH gene is localized outside the donor region, the presence of non-insect DNA in its vicinity strongly supports horizontal gene transfer (HGT) as the most parsimonious explanation.

### 3.6. Novel clock gene isoforms identified in Hemiptera

Gene presence or loss is not the only way to shape gene phenotypic output. We analyzed the splicing of mammalian-type *cryptochrome* (*m-cry*) and *period* (*per*) in several heteropteran species and focused on how it could affect the variability of the translated proteins. mCRY protein possesses two major domains, N-terminal DNA photolyase and C-terminal FAD binding 7 domain (Fig. 7). In all species analyzed, *N. lugens*, *P. apterus* and *H. halys*, *m-cry* is a subject of alternative splicing, giving rise to the truncated isoforms. *N. lugens m-cry* has two predicted transcripts, each transcribed from an alternative exon. Transcript X1 encodes the full protein with complete DNA Photolyase and FAD binding 7 domains. The shorter isoform X2 transcription starts from non-coding (5’UTR) exon 4 and lacks ∼68% of the Photolyase domain. Similarly, *H. halys* isoform X4 starts from an alternative exon 4, which leads to the lack of ∼67% of the Photolyase domain. A remarkable splicing pattern was found in *P. apterus* and *H. halys*, where we found *m-cry* isoforms with retained intron 10, splitting the FAB binding 7 domain-encoding exons (Fig. 7). Translation of retained-intron isoforms in both species leads to truncated mCRY proteins, with only ∼48% (*P. apterus*) and ∼22% (*H. halys*) fragment of the FAB binding 7 domain. The domain fragment in *H. halys* is below the threshold to be predicted as the FAD binding 7 domain by the PFAM database algorithm, and thus, the domain was not highlighted in Fig. 7. Interestingly, the ratio of intron-retained/full *m-cry* mRNA reads reached ∼32% in *P. apterus* Oxford Nanopore Technology brain transcriptomes. In *P. apterus*, C and D *per* isoforms are likely destroyed via NMD if translated from the first canonical AUG (isoforms C1 and D1) or translated into PER isoforms C2 and D2 if (downstream) alternative translation start site is preferred (Fig. 7). Whether initial codon skipping happens in vivo and gives rise to C2 and D2 isoforms is to be determined.

**Fig. 4.**
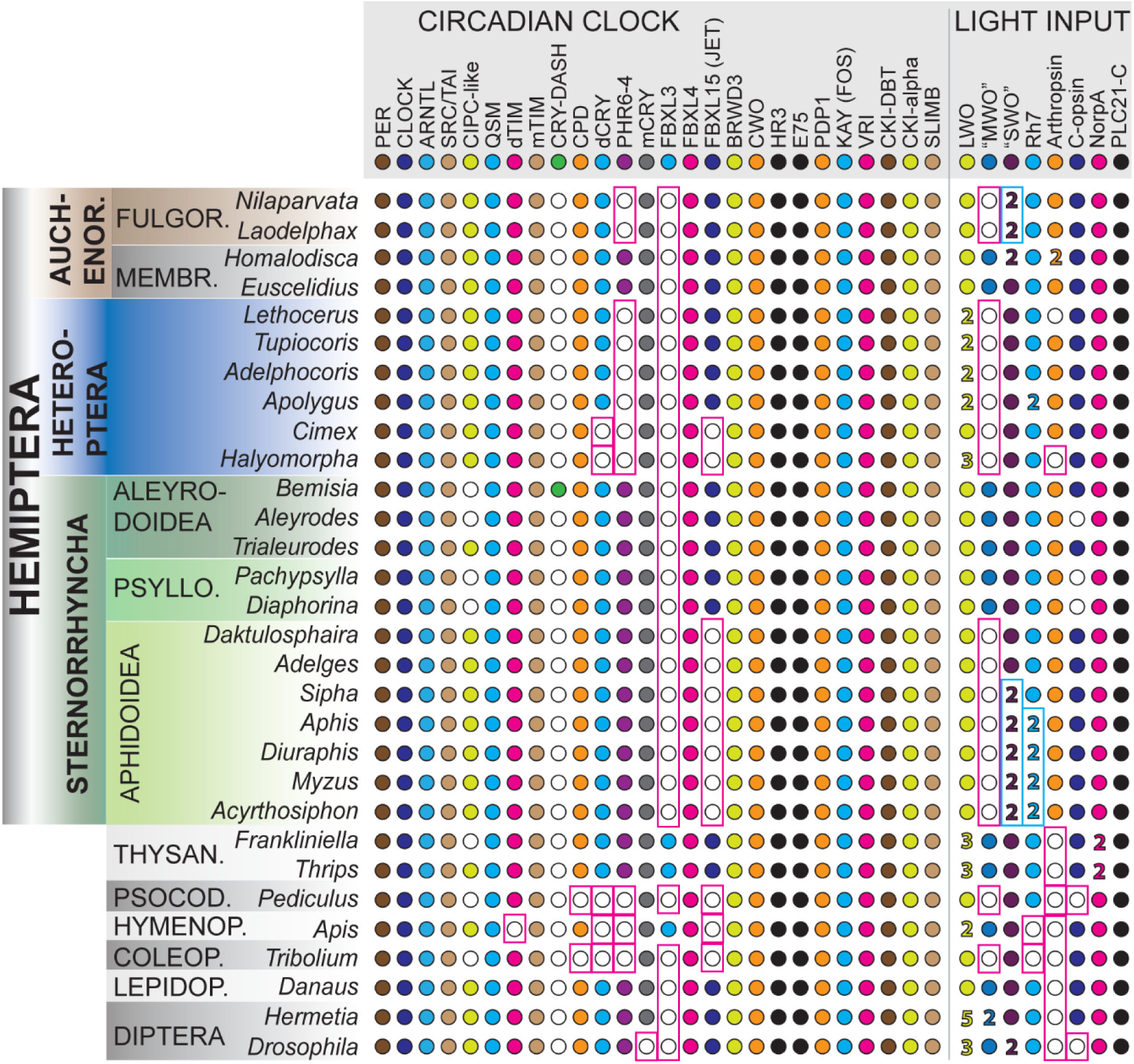
Inventory of circadian clock setup and light input components. Representative insect species are shown with gene presence (full circle) or absence (empty circle) indicated. Lineage-specific losses are highlighted with magenta rectangles, while lineage-specific duplications are highlighted with turquoise rectangles. Numbers represent the presence of multiple paralogs within a single taxon. Abbreviations are as follows: FULGOR. – Fulgoroidea, MEMBR. – Membracoidea, PSYLLO. –Psylloidea, THYSAN. –Thysanoptera, PSOCOD. –Psocodea, HYMENOP. –Hymenoptera, COLEOP. –Coleoptera, LEPIDOP. –Lepidoptera.

**Fig. 5.**
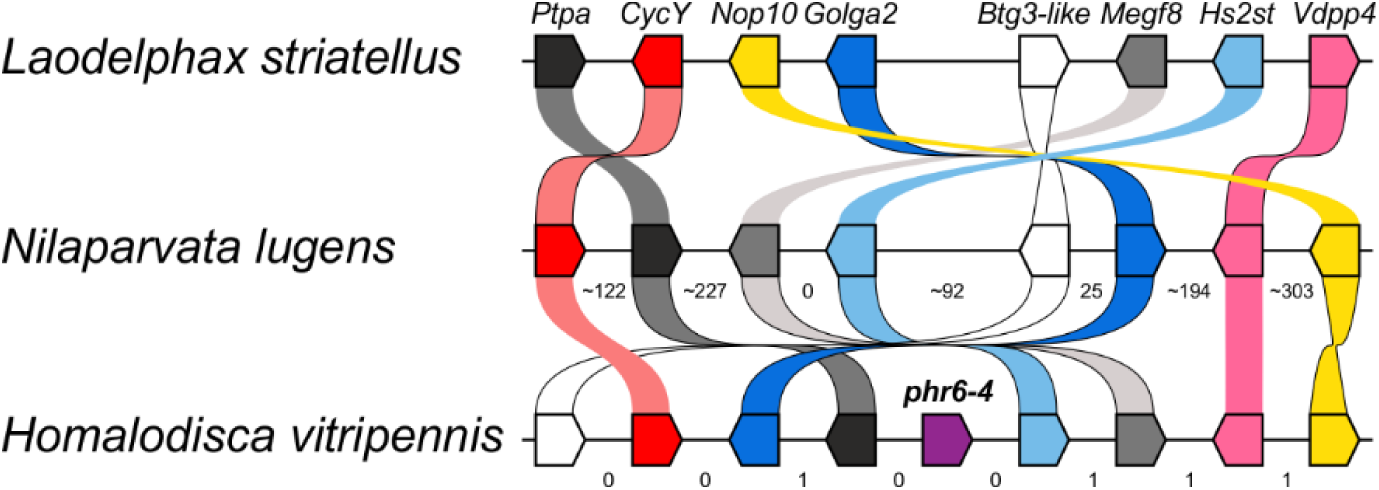
Gene synteny supports the loss of *(6-4)-photolyase* (*phr 6-4*) in *Nilaparvata lugens* and other Fulgoroidea. *Laodelphax striatellus* and *N. lugens* (both Fulgoroidea) have lost *phr6-4*, which remains present in the sister family Membracoidea, as represented by *Homalodisca vitripennis*. The *phr6-4* upstream and downstream syntenic genes from *H. vitripennis* are scattered across a single contig or scaffold in the genomes of *N. lugens* and *L. striatellus*. Syntenic genes are color-coded for easier tracking. Numbers between genes in *N. lugens* and *H. vitripennis* indicate the number of protein-coding genes separating the respective syntenic genes. Note the significant reorganization of the presented genes within the contigs.

**Fig. 6.**
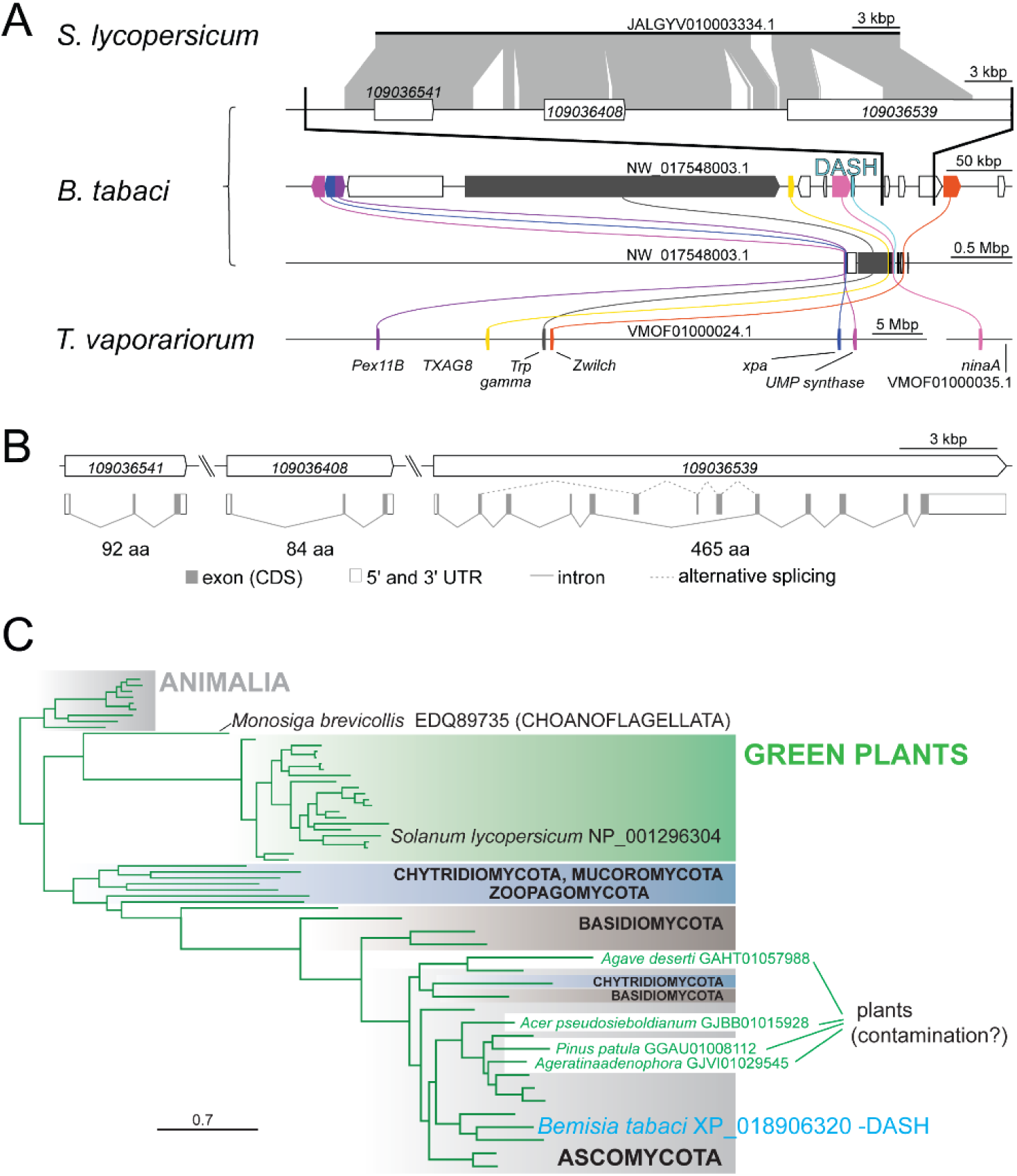
*Bemisia tabaci cryptochrome-DASH* locus analysis supports horizontal gene transfer. (A) Seven protein-coding genes located upstream and downstream of the *B. tabaci cryptochrome-DASH* were also found in the related whitefly *Trialeurodes vaporariorum*; however, the gene order is significantly rearranged. These syntenic genes are color-coded for easier tracking (note that each sequence is presented at a different scale). Interestingly, a BLAST search of the *B. tabaci cry-DASH* locus DNA sequence against GenBank genome databases revealed ∼90% identity between a 33-kbp genomic fragment of the tomato plant (*Solanum lycopersicum*; JALGYV010003334.1) and a ∼40-kbp segment of the *B. tabaci* genomic sequence (NW_017548003.1) located downstream of the *cry-DASH* gene. (B) Details of three genes predicted in the locus downstream of *cry-DASH*. Note that the protein-coding sequence constitutes only a small portion of each predicted hypothetical gene. (C) Phylogenetic analysis of CRY-DASH proteins. Animal CRY-DASH proteins served as an outgroup (top, Animalia). One cluster contains only CRY-DASH from green plants (green background, middle part). The bottom branch includes several representatives of fungal lineages (Chytridiomycota, Mucoromycota, Zoopagomycota, Basidiomycota, and Ascomycota), highlighted with different background colors. The *Bemisia* cry-DASH branches within *Ascomycota*. Four plant “CRY-DASH” protein sequences branching within fungi were identified in the Transcriptome Shotgun Assembly (TSA) databases. These sequences are highly suspicious and are likely the result of cross-contamination of the plant sample with fungi.

**Fig. 7.**
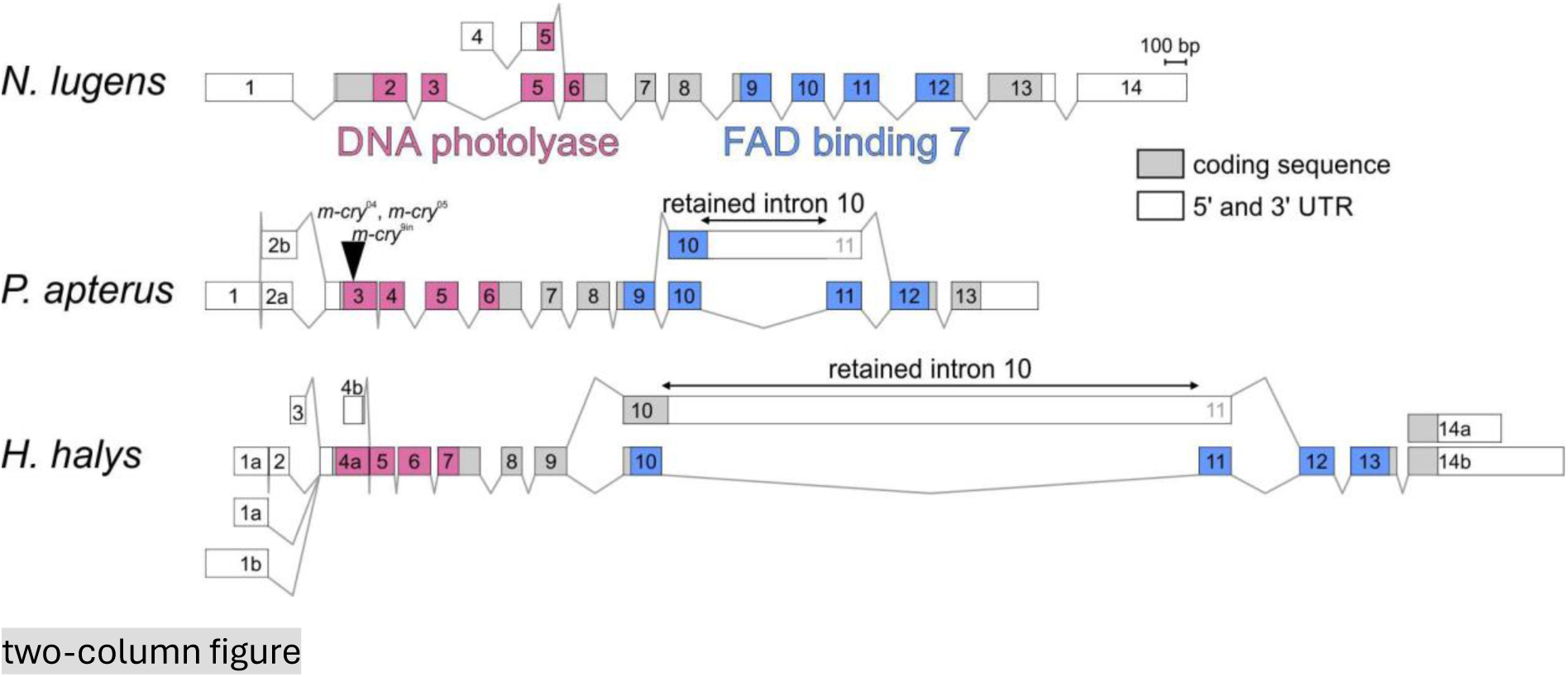
mCRY splicing isoforms. The *m-cry* gene of *N. lugens*, *P. apterus*, and *H. halys* is subject to alternative splicing, which generates some truncated mCRY protein isoforms. In *N. lugens*, transcription from an alternative exon 4 produces an mCRY isoform with a significantly shorter DNA photolyase domain. Retained intron 10 in *P. apterus* and *H. halys* introduces a premature stop codon, truncating the coding sequence of the mCRY FAD-binding 7 domain. These shorter mCRY proteins are either non-functional or targeted for transcript suppression via nonsense-mediated mRNA decay. All exons are shown to scale and numbered in the 5’ → 3’ direction, with letters depicting exon sub-variants. The black arrowhead marks the positions of *P. apterus m-cry* mutants (Kotwica-Rolinska et al., 2022).

A complex splicing pattern was found in the 5’ region of the *period* gene in the analyzed species, giving rise to PER variants differing in the N-terminal part. Reference *D. melanogaster period* is not alternatively spliced in this gene region. The only translated PER protein isoform possesses an N-terminal nuclear localization signal (NLS, Chang and Reppert 2003), two PAS domains, mid-protein NLS (Fig. 8), and several phosphocluster regions, which are phosphorylated to ensure PER stability, inhibitory potency and modulation of temperature compensation phenotype, forming thus a phospho-switch to balance diverse PER functions throughout the day-night cycle (Joshi et al., 2022). One of the phosphoclusters, the N-terminal phosphodegron, is responsible for PER degradation in the proteasome (Chiu et al., 2008; Kivimäe et al., 2008). It includes Serine 44,45 and 47 (numbering according to *D. melanogaster* PER, Chiu et al., 2008). S47 is phosphorylated by DBT and together with nearby phosphorylated sites, phospho-S47 generates a high-affinity atypical SLIMB-binding site (Chiu et al., 2008). The phosphodegron with flanking sequences is conserved in full PER protein isoforms in analyzed species, PER isoform X1 in *N. lugens*, isoform A in *P. apterus* and isoform X1 in *H. halys* (Fig. 8B). Unlike *D. melanogaster*, analyzed heteropteran PERs contain an additional C-terminal Period circadian-like domain (PeriodC domain, PFAM ID: IPR022728) (Fig. 8A). The full PER represents only a subset of possible splice variants in analyzed hemipteran species, with several gradually truncated PERs. In *N. lugens* and *H. halys*, transcription from alternative exons 9 and 7 leads to an expression of X19 and X8 isoforms, respectively, missing the phosphodegron and N-terminal NLS. Most intriguing *per* isoforms, generated by the alternative splicing, were detected in *P. apterus* and *H. halys*. Transcription of *P. apterus per* isoform B from exon 2 and *H. halys per* isoforms X6 (and X7) from exon 3 alternative transcription start site reveals the first initiation codon (AUG) in exon 4 but in reading frame 2 (depicted by a red line in Fig. 8A). The same region is in full-PER isoforms read in frame 1. In both species, translation starts with exon 4 and the translation in reading frame 2 continues through exon 6 (*P. apterus*) or 5 (*H. halys*). The absence of exon 7 (*P. apterus*) or exon 6 (*H. halys*) in the ‘alternative’ B and X6/X7 isoforms, respectively, brings the rest of the open reading frame into line with reading frame 1, thus avoiding premature stop codon and enabling translation of the downstream canonical PER sequence and domains (PAS, mid-protein NLS, PeriodC). Remarkably, the exons with dual/overlapping reading frames encode NLS in both frames (!), although the exact AA composition differs. *P. apterus* full-length mRNA Oxford Nanopore Technology (ONT) reads further revealing the presence of minor versions of transcripts encoding isoforms A and B, in Fig. 8 presented as isoforms C and D. Those isoforms either lack (isoform C) or contain (isoform D) frame-adjusting exon 7, thus decoupling the frames.

**Fig. 8.**
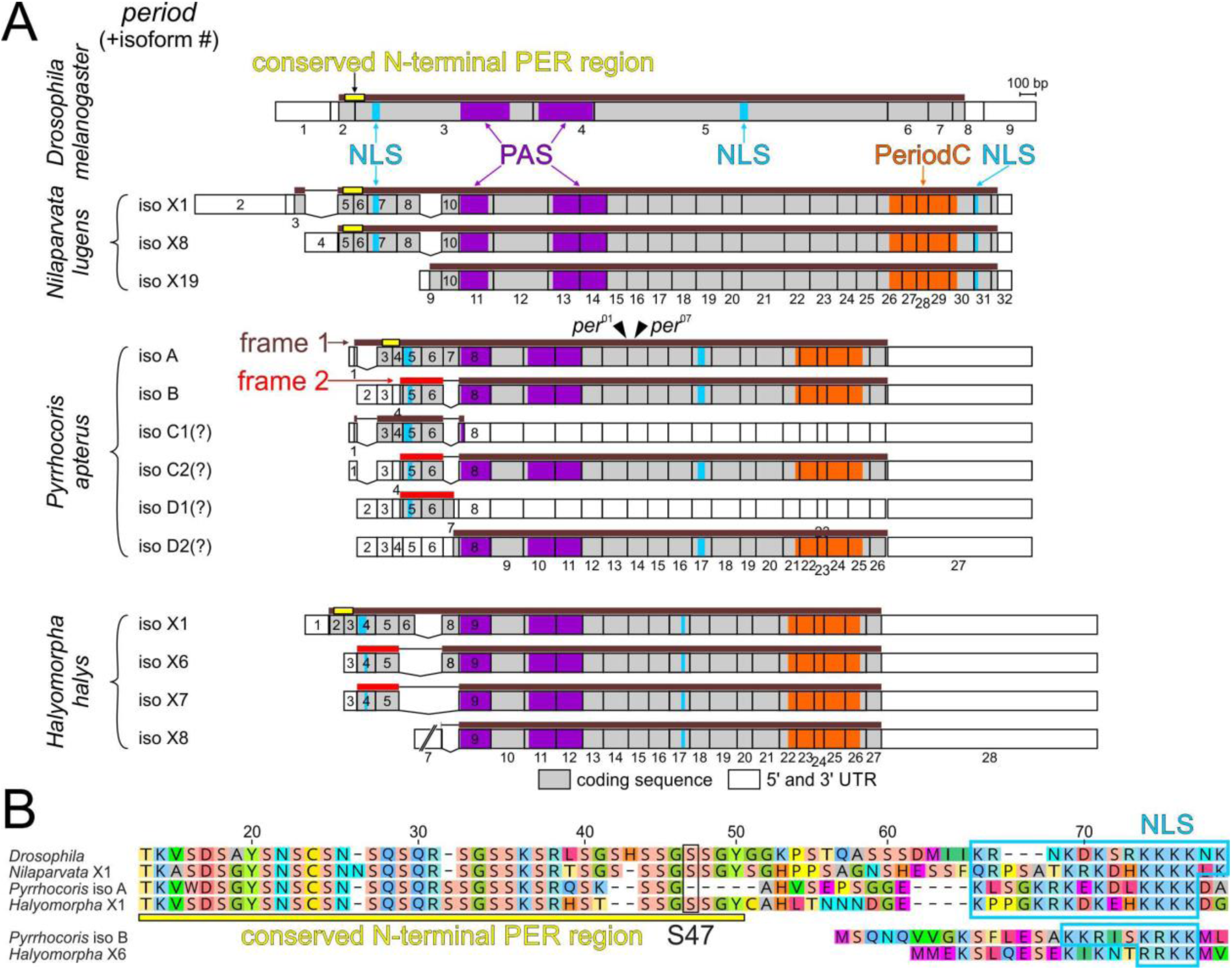
PERIOD N-terminal isoforms are generated by alternative splicing and dual-reading-frame translation of specific exons. (A) Full-length PER proteins are translated in reading frame 1 (brown line) and possess conserved 32-33-amino-acid-long N-terminal PER region (yellow line). A gradient of N-terminally truncated PER is generated by alternative splicing in *N. lugens*, *P. apterus* and *H. halys*, but not *Drosophila melanogaster*. Some isoforms lack conserved N-terminal PER region, some lack N-terminal nuclear localization signal (NLS). *N. lugens per* isoforms are generated by transcription from alternative transcriptional start sites. Transcription of *P. apterus per* isoform B and *H. halys per* isoforms X6 and X7 from alternative transcription start sites leads to translation in reading frame 2 (red line). The absence of exon 7 (*P. apterus*) or exon 6 (*H. halys*) in the isoforms changes the reading frame back to frame 1, enabling the presence of the canonical PER domains (PAS, mid-protein NLS, PeriodC). Interestingly, dual/overlapping reading frames exons encode NLS in both frames. *P. apterus per* isoform C2 is likely destroyed via nonsense-mediated mRNA decay if translated from the canonical AUG or results into isoform C1 if (downstream) alternative translation start site is preferred. All exons are numbered in a 5’->3’ direction and presented in scale, except *H. halys* exon 7, which was shortened for space reasons. The black arrowhead depicts the positions of *P. apterus per* mutants (Kotwica-Rolinska et al., 2022a). See Methods and Materials for a detailed description of the protein domain and motifs’ prediction. (B) The alignment of conserved N-terminal PER region. The black rectangle highlights Serine 47 (S47), targeted in *Drosophila melanogaster* by DBT and upon phosphorylation targeting PER for degradation (Chiu et al., 2008). The yellow line depicts the conserved N-terminal PER region and the blue curved line predicted NLS.

## 4. Discussion

This study focused on circadian clock genes and opsins in Hemiptera. In principle, this is a gene inventory aimed at shedding light on the evolution of this time-measuring molecular device. While some earlier studies focused on individual species, such as the bean bug *Riptortus pedestris* (Ikeno et al., 2008) or the pea aphid *Acyrthosiphon pisum* (Cortés et al., 2010; Barberà et al., 2017), systematic comparisons, to our knowledge, have not been provided. Our earlier publication (Kotwica-Rolinska et al., 2022a) addressed the function of the majority of clock genes in the linden bug *Pyrrhocoris apterus*; however, the comparative part was limited to PER, dTIM, mTIM, cryptochromes, JET, and FBXL3 proteins. In addition to Hemiptera, our current description includes several reference/outgroup insect lineages. Thus, we can further expand the concept of what the ancestral type of insect circadian clock consisted of. The combination of mCRY and dCRY, typical of the lepidopteran clock, is further complemented by FBXL3 in Thysanoptera, suggesting that, at least at present, thrips possess the most complete circadian gene setup among insects. The reference insect species illustrate the diversity of clock setups in insects, including the notable loss of dCRY, (6-4) photolyase, and JET. In the case of *Apis*, the clock setup is further reduced by the loss of dTIM. The comparison of *Drosophila* and *Hermetia*, both members of Diptera, illustrates the mCRY loss unique to Cyclorrhapha.

Although some important differences in the mechanism of the circadian clock exist, such as the involvement of mCRY in numerous species contrasting with *Drosophila* (Ikeno et al., 2011a; Zhang et al., 2017; Werckenthin et al., 2020) and the altered function of dTIM (Kamae and Tomioka, 2012; Kotwica-Rolinska et al., 2022a), it is still feasible to extrapolate clock properties based on the *Drosophila* mechanisms. For example, the loss of dCRY and JET in *P. apterus* fits with the lower sensitivity to light for entrainment (Kaniewska et al., 2020). In aphids, dCRY is unexpectedly stable upon constant light illumination (Colizzi et al., 2021). This stability can be explained by the loss of JET and further correlates with the accelerated evolution of dTIM in aphids and phylloxera (Bullo et al., 2024). Whether dCRY participates in the light entrainment of the circadian clock or plays some role in the photoperiodic timer in aphids, remains elusive.

However, the light input to the clock is rather complex when parallel dCRY-independent pathways have been reported. In some cases, it is not clear whether they represent a conserved mechanism or are mere species-specific idiosyncrasies. For example, the role of QSM in light responsiveness has not been explored beyond *Drosophila melanogaster*. In the case of opsins, their role in the circadian clock is well-established in several insect species. However, the presence of multiple opsin paralogs makes their research technically challenging. For example, seven opsin genes are recognized in *Drosophila*. Thus, despite the remarkable genetic tools available to this model organism, even combining all available mutations into one strain becomes a non-trivial task. For example, the role of Rh-7 is still being uncovered (Senthilan and Helfrich-Förster 2016).

A second research challenge is the spectral overlap in the sensitivity of several opsins. Importantly, the spectral sensitivity has been experimentally determined for only a minimal number of opsins, whereas the properties of many others remain elusive. Therefore, the definition of particular groups in this study should be understood based on their evolutionary position rather than their actual light-detection properties. Notably, some experimental evidence highlights specific residues that are key for determining wavelength sensitivity. When these residues are mutated, spectral shifts occur, as suggested for one of Aphidoidea (and possibly for Fulgoroidea) SWO paralogs (Collantes-Alegre et al., 2018). Such light-sensing adaptability would contribute to physiological plasticity and potentially be a simple solution compensating for MWO gene loss.

In addition to gene losses, we identified a unique horizontal gene transfer in *B. tabaci*, where a fungal *cry-DASH* had been inserted into the insect genome. The presence of an exogenous DNA in the vicinity of the *cry-DASH* locus in the *B. tabaci* genome supports HGT as a feasible origin of *cry-DASH* in only one insect species. Our analysis shows that the *Solanum lycopersicum* genomic fragment is mapped in several neighboring regions, suggesting ongoing whitefly-specific evolution of the locus. Although *S. lycopersicum* seems to be the donor of the HGT sequence, we cannot rule out the possibility that the true donor species has not been sequenced yet and *S. lycopersicum* genomic fragment is just the closest species present in the GenBank. Phylogenetic analysis of CRY-DASH proteins questioned the plant origin of *B. tabaci* CRY-DASH, placing it into a ‘fungal-specific’ cluster. Indeed, *B. tabaci* CRY-DASH is more related to fungal than plant CRY-DASH proteins, including CRY-DASH from *S. lycopersicum* genome. Whether the physical vicinity of *B. tabaci cry-DASH* locus and exogenous DNA represents a single or two independent HGT events is unclear. The gene synteny comparison with the related *Trialeurodes vaporariorum* whitefly genome underpins major genomic rearrangements, further masking the origin of the *cry-DASH* locus. In any case, *B. tabaci* is a polyphagous species with ∼500 host plants in its repertoire, thus facing many challenges when feeding upon different hosts. Over 49 genes were likely obtained from plant hosts via HGT by *B. tabaci* (Lapadula et al., 2020; Xia et al., 2021; Gilbert and Maumus 2022), many linked to insect-plant interactions. The HGT can provide an alternative for gaining new capabilities through classical gene duplication and evolution. Although the *cry-DASH* in *B. tabaci* is transcribed, whether possessing this gene constitutes an advantage remains to be determined.

The transcriptome analysis revealed a remarkable dual reading frame in *P. apterus* and *H. halys period* transcripts. The Serine-rich phosphodegron is well conserved in full-length PER proteins in all four analyzed species, with *P. apterus* PER lacking Ser47 (replaced by Ala50). At least one isoform lacking PER phosphodegron is annotated in *N. lugens*, *P. apterus*, and *H. halys*, probably omitting DBT– and SLIMB-specific regulation in the N-terminal region (Fig. 8A), however, specific point mutations in *D. melanogaster* PER phosphodegron keep downstream parts of the protein still accessible to DBT and SLIMB regulation (Chiu et al., 2008).

Unlike *Drosophila*, where PER heterodimerizes with and is stabilized by dTIM, all analyzed species possess mCRY (Fig. 3 and Kotwica-Rolinska et al., 2022a), a typical mammalian PER partner. We can speculate if phosphodegron-less PERs represent less SLIMB-sensitive isoforms or denote the variability of PER heterodimerization with dTIM and mCRY. The presence of phosphodegron-less PER isoforms in *P. apterus* and *H. halys* suggests that the lack of phosphodegron outweighs the AA variability in N-terminal NLS, encoded by the dual-frame exons, and a necessity to bring those exons in frame with the rest of the PER protein to contain PAS and other downstream domains. Manipulation of N-terminal and mid-PER NLS sequences in *D. melanogaster* PER suggested the mid-PER NLS is the main nuclear localization signal, however, N-terminal NLS still participates in PER nuclear import (Chang and Reppert, 2003). Thus, an ‘alternative’ NLS with a different AA composition might still be sufficient to complement mid-PER NLS.

Alternative splicing serves as yet another layer for controlling clock gene expression. For instance, retention of an intron in the central part of the *d-tim* gene in *Drosophila* results in transcripts that cannot encode a stable dTIM protein (Martin Anduaga et al., 2019; Foley et al., 2019; Shakhmantsir et al., 2018). Furthermore, these mRNAs contain premature stop codons and are therefore likely destroyed by NMD. The high frequency of intron retention in the *m-cry* transcripts of *P. apterus* invites speculation that a similar regulatory mechanism might be at play. It will be interesting to determine whether similar intron retention occurs in other species. Since the stability of RNA-RNA interactions is temperature-dependent, alternative splicing is often influenced by temperature. Indeed, the temperature-dependent splicing of *d-tim* in drosophilid flies likely serves as a regulatory mechanism. Whether *m-cry* splicing is affected by temperature remains unknown, as does its potential regulatory role in the circadian clock. However, because the mCRY protein is a key clock component in numerous species, including *P. apterus*, significant regulation of its levels could, at least theoretically, have a profound impact on clock function.

What is responsible for the observed genetic flexibility of clock setups? As these time-measuring devices are supposed to perform a conserved task, which is to keep ticking with a period of 24 hours, one would assume that the genetic components should be very conserved. While the general mechanism of the transcriptional/translational feedback loop is found in both deuterostomes (including mice, humans, and zebrafish) and protostomes (insects, mollusks, and annelids), the participating players are conserved only partially. Either gene losses or a high degree of sequence variability in the protein sequences are observed. One possible mechanism allowing for gene loss and rapid evolution of some components is the functional redundancy of the system. For example, the light input to the clock is achieved by several pathways; thus, alteration of one of them does not result in complete “clock blindness.” Therefore, losing one of these pathways only impacts sensitivity, but the organism still perceives particular stimuli. This change in sensitivity might even be beneficial. For example, insects have colonized various environments that differ in photoperiod (day-to-night length). In the most extreme latitudes, nearly constant light is available during summer. Under such conditions, lower light sensitivity might be an advantage. Indeed, this is the case for *Drosophila ezoana*, which is partially rhythmic even under constant light conditions. Furthermore, this and some other species from high latitudes, such as *Chymomyza costata*, are not rhythmic in constant darkness (Bertolini et al., 2019). In the latter species, a unique recessive mutation in the *d-tim* gene is associated with a malfunction in photoperiodic timing and impacts the expression of additional clock genes (Pavelka et al., 2003; Stehlik et al., 2008; Kobelkova et al., 2010). The possible connection between photoperiodic timers and the circadian clock has been long discussed (Kostal 2011; Dolezel 2015; Saunders 2005, 2010; Bradshaw and Holzapfel, 2010). The recruitment of circadian clock components by the photoperiodic timer seems to be a logical explanation for establishing a relatively complex device measuring photoperiod from already available “tools”. Indeed, this view is further supported by the participation of several clock genes and circadian neuropeptide PDF in the photoperiodic timer in several species of Hemiptera (Ikeno et al., 2010; Ikeno et al., 2011b, 2011c, 2013; Urbanova et al., 2016; Kotwica-Rolinska et al., 2017, 2022b; Kaniewska et al., 2024). However, although the evidence is impressive, the actual involvement of the entire circadian clock machinery in the photoperiodic timer has not been explored and the connection between the two devices is more complicated. One important critique is that only some components might have been recruited (Emerson et al., 2009). Gene pleiotropy, such as when circadian clock components in the insect gut interact in a non-canonical way, suggests that the connection between the circadian clock and seasonality is indeed more complex (Bajgar et al., 2013a, 2013b; Dolezel et al., 2007).

The circadian clock genes also define the activity phase, and this role might have shaped their evolution. In this regard, notable geographic variability in the free-running period has been reported for the linden bug *P. apterus*, for the wasp *Nasonia vitripennis*, and in the eclosion rhythm of *Drosophila littoralis* and *D. subobscura* (Pivarciova et al., 2016; Paolucci et al., 2019; Lankinen, 1986; Lankinen, 1993).

Another key role of the circadian clock is related to navigation during long migrations. The position of the Sun serves as a suitable orientation cue as long as the organism can compensate for its positional change during the day. The time-compensated sun compass is intensively studied in the iconic monarch butterfly, *Danaus plexippus*, in which the ability to detect the magnetic field contributes to migration either by fine-tuning the navigation system or serving as a backup mechanism (Merlin et al., 2020). Moreover, the role of these components is further complicated by the participation of cryptochromes in the ability to detect the magnetic field and relay this information. The interference of the magnetic field with the effects of light has been well documented in *Drosophila* and the German cockroach *Blattella germanica* (Fedele et al., 2014a, 2014b; Yoshii et al., 2009). In the case of *Drosophila*, the response requires dCRY. The strongest phenotypes pointing to the role of dCRY in directional magnetoreception have been observed in the monarch butterfly (Wan et al., 2021), although, in some species, mCRY has been suggested (Bazalova et al., 2016; Netusil et al., 2021). Interestingly, a change in geomagnetic field intensity alters migration-associated traits in *Nilaparvata* (Wan et al., 2020).

The complex involvement of the circadian clock genes in various time-measuring devices, navigation, magnetoreception, and seasonality provides interesting scientific problems but may also be connected to applied research. Numerous hemipteran species are important pests and understanding the genetic basis of their daily activity preference, seasonal calendar, and navigation can be practical. Similarly, our analysis points to the importance of alternative splicing isoforms, such as the non-canonical PER variants and intron retention in *m-cry*. In both cases, and many others found in the future, it will also be interesting to explore whether the alternative splicing is affected by temperature. Perhaps, some alternative splicing events even participate in temperature compensation mechanisms or contribute to the regulation of temperature-dependent daily activity patterns. Therefore, it is important to identify and annotate all predicted gene isoforms when dealing with genomic or transcriptomic data. As recently shown by Chikhaoui et al. (2024), the loss of PER proteins in the liver of *mPer1*/*mPer2* double mutant mice seems to directly alter alternative splicing likely through the mis-localization of serine/arginine-rich splicing factors (SRSF) within the nucleus. Thus, alternative splicing might represent still rather undervalued level of clock regulatory mechanism.

While the *in silico* predictions point to important changes and irregularities in the clock setup, functional analysis is key for identifying underlying mechanisms. In this regard, stable genomic modifications are the most powerful approach to entering hemipteran research including challenging groups such as aphids (Xue et al., 2018; Kotwica-Rolinska et al., 2019; Reding and Pick, 2020; Le Trionnaire et al., 2019; de Souza Pacheco et al., 2022). To make gene editing accessible to a broader range of non-model species, a more versatile means of Cas9/gRNA delivery are emerging (Mocchetti et al., 2024; Shirai et al., 2022) and less harmful DNA-editing enzymes are tested (Thakkar et al., 2023; Doll et al., 2023; Li et al., 2018). Therefore, we can assume that some significant progress will be made in understanding the circadian clock genes in either interesting biological problems or in agronomically important pest species.

## Author contribution

Conceptualization: D.D., V.S., Data curation: V.S., H.T., D.D., Formal analysis: D.D., V.S., H.T., Funding acquisition: D.D., Investigation: D.D., V.S., H.T., Methodology: D.D., V.S., Resources: D.D., V.S., Supervision: D.D., V.S., Validation: D.D., V.S., Visualization: D.D., V.S., Writing – original draft: D.D., V.S., Writing – review and editing: D.D., V.S.

## Appendix A. Supplementary data

**Fig. S1A.** Protein alignment of selected opsins from the Rh7, SWO, and MWO clades.

**Fig. S2** Phylogenetic analysis of CRY-DASH proteins.

**Table S1.** Synteny in *6-4 photolyase* locus

**Table S2.** *Cryptochrome-DASH* synteny

**Table 1.** An enhanced list of issues to focus on if studying clock genes in heteropteran insects.

|  |  |
| --- | --- |
| gene search | • genomic and transcriptomic databases<br>• presence/absence and identity<br>• duplications & losses<br>• horizontal gene transfer |
| gene expression | • alternative promoters & protein isoforms<br>• alternative UTRs & retained introns<br>• nonsense-mediated mRNA decay<br>• transcription factors & enhancers<br>• upstream open reading frames (uORFs)<br>• microRNAs |
| phenotypic properties | • locomotor activity in constant darkness<br>• impact of constant light on rhythmicity (particularly comparison of lineages with/without dCRY)<br>• entrainment cues – lineages with/without dCRY<br>• migration, seasonality |
| reverse genetics | • RNA interference & genome editing (CRISPR/Cas9)<br>• phenotypes (locomotor activity, gene expression, rhythmicity, period) |

## Supporting information

upplementary material:

## References

1. Allada, R., White, N.E., So, W.V., Hall, J.C., Rosbash, M., 1998. A mutant Drosophila homolog of mammalian Clock disrupts circadian rhythms and transcription of period and timeless. Cell 93, 791–804. doi: 10.1016/s0092-8674(00)81440-3

2. Bai, J., Uehara, Y., Montell, D.J., 2000. Regulation of invasive cell behavior by taiman, a Drosophila protein related to AIB1, a steroid receptor coactivator amplified in breast cancer. Cell 103, 1047–1058. doi: 10.1016/s0092-8674(00)00208-7

3. Bajgar, A., Dolezel, D., Hodkova, M., 2013a. Endocrine regulation of non-circadian behavior of circadian genes in insect gut. J Insect Physiol 59, 881–886. doi: 10.1016/j.jinsphys.2013.06.004

4. Bajgar, A., Jindra, M., Dolezel, D., 2013b. Autonomous regulation of the insect gut by circadian genes acting downstream of juvenile hormone signaling. Proc Natl Acad Sci U S A 110, 4416–4421. doi: 10.1073/pnas.1217060110

5. Barberà, M., Collantes-Alegre, J.M., Martinez-Torres, D., 2017. Characterisation, analysis of expression and localisation of circadian clock genes from the perspective of photoperiodism in the aphid Acyrthosiphon pisum. Insect Biochem Mol Biol 83, 54–67. doi: 10.1016/j.ibmb.2017.02.006

6. Barberà, M., Collantes-Alegre, J.M., Martinez-Torres, D., 2022. Mapping and quantification of cryptochrome expression in the brain of the pea aphid Acyrthosiphon pisum. Insect Mol Biol 31, 159–169. doi: 10.1111/imb.12747

7. Bazalova, O., Kvicalova, M., Valkova, T., Slaby, P., Bartos, P., Netusil, R., Tomanova, K., Braeunig, P., Lee, H.J., Sauman, I., Damulewicz, M., Provaznik, J., Pokorny, R., Dolezel, D., Vacha, M., 2016. Cryptochrome 2 mediates directional magnetoreception in cockroaches. Proc Natl Acad Sci U S A 113, 1660–1665. doi: 10.1073/pnas.1518622113

8. Bertolini, E., Schubert, F.K., Zanini, D., Sehadova, H., Helfrich-Forster, C., Menegazzi, P., 2019. Life at High Latitudes Does Not Require Circadian Behavioral Rhythmicity under Constant Darkness. Curr Biol 29, 3928–3936. doi: 10.1016/j.cub.2019.09.032

9. Blau, J., Young, M.W., 1999. Cycling vrille expression is required for a functional Drosophila clock. Cell 99, 661–671. doi: 10.1016/s0092-8674(00)81554-8

10. Bradshaw, W.E., Holzapfel, C.M., 2010. What Season Is It Anyway? Circadian Tracking vs. Photoperiodic Anticipation in Insects. J Biol Rhythms 25, 155–165. doi: 10.1177/0748730410365656

11. Bullo, E., Chen, P., Fiala, I., Smykal, V., Dolezel D., 2024. Coevolution of Drosophila-type Timeless with Partner Clock Proteins. bioRxiv. doi: 10.1101/2024.12.25.628932

12. Ceriani, M.F., Darlington, T.K., Staknis, D., Mas, P., Petti, A.A., Weitz, C.J., Kay, S.A., 1999. Light-dependent sequestration of TIMELESS by CRYPTOCHROME. Science 285, 553–556. doi: 10.1126/science.285.5427.553

13. Chang, D.C., Reppert, S.M., 2003. A novel C-terminal domain of drosophila PERIOD inhibits dCLOCK:CYCLE-mediated transcription. Curr Biol 13, 758–762. doi: 10.1016/s0960-9822(03)00286-0

14. Chen, K.F., Peschel, N., Zavodska, R., Sehadova, H., Stanewsky, R., 2011. QUASIMODO, a Novel GPI-Anchored Zona Pellucida Protein Involved in Light Input to the Drosophila Circadian Clock. Curr Biol 21, 719–729. doi: 10.1016/j.cub.2011.03.049

15. Chikhaoui, L., Mamgain, K., Seki, M., Blanco, C., Sassolas, F., Folco, E., Sery, D., Suzuki, Y., Ananthasubramaniam, B., Padmanabhan, K., 2024. Circadian PERIOD proteins sculpt the mammalian alternative splicing landscape. bioRxiv doi: 10.1101/2024.12.23.630108

16. Chiu, J.C., Vanselow, J.T., Kramer, A., Edery, I., 2008. The phospho-occupancy of an atypical SLIMB-binding site on PERIOD that is phosphorylated by DOUBLETIME controls the pace of the clock. Genes Dev 22, 1758–1772. doi: 10.1101/gad.1682708

17. Colizzi, F.S., Beer, K., Cuti, P., Deppisch, P., Martinez Torres, D., Yoshii, T., Helfrich-Forster, C., 2021. Antibodies Against the Clock Proteins Period and Cryptochrome Reveal the Neuronal Organization of the Circadian Clock in the Pea Aphid. Front Physiol 12, 705048. doi: 10.3389/fphys.2021.705048

18. Collins, B., Mazzoni, E.O., Stanewsky, R., Blau, J., 2006. Drosophila CRYPTOCHROME is a circadian transcriptional repressor. Curr Biol 16, 441–449. doi: 10.1016/j.cub.2006.01.034

19. Collantes-Alegre, J.M., Mattenberger, F., Barberà, M., Martínez-Torres, D., 2018. Characterisation, analysis of expression and localisation of the opsin gene repertoire from the perspective of photoperiodism in the aphid Acyrthosiphon pisum. J Insect Physiol 104, 48–59. doi: 10.1016/j.jinsphys.2017.11.009

20. Cortés, T., Ortiz-Rivas, B., Martinez-Torres, D., 2010. Identification and characterization of circadian clock genes in the pea aphid Acyrthosiphon pisum. Insect Mol Biol 19, 123–139. doi: 10.1111/j.1365-2583.2009.00931.x

21. Cyran, S.A., Buchsbaum, A.M., Reddy, K.L., Lin, M.C., Glossop, N.R.J., Hardin, P.E., Young, M.W., Storti, R.V., Blau, J., 2003. vrille, Pdp1, and dClock form a second feedback loop in the Drosophila circadian clock. Cell 112, 329–341. doi: 10.1016/s0092-8674(03)00074-6

22. D’Costa, A., Reifegerste, R., Sierra, S., Moses, K., 2006. The Drosophila ramshackle gene encodes a chromatin-associated protein required for cell morphology in the developing eye. Mech Dev 123, 591–604. doi: 10.1016/j.mod.2006.06.007

23. de Souza Pacheco, I., Doss, A.A., Vindiola, B.G., Brown, D.J., Ettinger, C.L., Stajich, J.E., Redak, R.A., Walling, L.L., Atkinson, P.W., 2022. Efficient CRISPR/Cas9-mediated genome modification of the glassy-winged sharpshooter Homalodisca vitripennis (Germar). Sci Rep 12, 6428. doi: 10.1038/s41598-022-09990-4

24. DeOliveira, C.C., Crane, B.R., 2024. A structural decryption of cryptochromes. Front Chem 12, 1436322. doi: 10.3389/fchem.2024.1436322

25. Dolezel, D., 2015. Photoperiodic time measurement in insects. Curr Opin Insect Sci 7, 98–103. doi: 10.1016/j.cois.2014.12.002

26. Dolezel, D., 2023. Molecular Mechanism of the Circadian Clock. In: Numata, H., Tomioka, K. (eds) Insect Chronobiology. Entomology Monographs. Springer, Singapore. doi: 10.1007/978-981-99-0726-7_4

27. Dolezel, D., Sauman, I., Kost’al, V., Hodkova, M., 2007. Photoperiodic and food signals control expression pattern of the clock gene, period, in the linden bug, Pyrrhocoris apterus. J Biol Rhythms 22, 335–342. doi: 10.1177/0748730407303624

28. Dolezelova, E., Dolezel, D., Hall, J.C., 2007. Rhythm defects caused by newly engineered null mutations in Drosophila’s cryptochrome gene. Genetics 177, 329–345. doi: 10.1534/genetics.107.076513

29. Doll, R.M., Boutros, M., Port, F., 2023. A temperature-tolerant CRISPR base editor mediates highly efficient and precise gene editing in Drosophila. Sci Adv 9, eadj1568. doi: 10.1126/sciadv.adj1568

30. Emerson, K.J., Bradshaw, W.E., Holzapfel, C.M., 2009. Complications of complexity: integrating environmental, genetic and hormonal control of insect diapause. Trends Genet 25, 217–225. doi: 10.1016/j.tig.2009.03.009

31. Emery, P., Stanewsky, R., Helfrich-Forster, C., Emery-Le, M., Hall, J.C., Rosbash, M., 2000. Drosophila CRY is a deep brain circadian photoreceptor. Neuron 26, 493–504. doi: 10.1016/s0896-6273(00)81181-2

32. Fedele, G., Edwards, M.D., Bhutani, S., Hares, J.M., Murbach, M., Green, E.W., Dissel, S., Hastings, M.H., Rosato, E., Kyriacou, C.P., 2014a. Genetic analysis of circadian responses to low frequency electromagnetic fields in Drosophila melanogaster. PLoS Genet 10, e1004804. doi: 10.1371/journal.pgen.1004804

33. Fedele, G., Green, E.W., Rosato, E., Kyriacou, C.P., 2014b. An electromagnetic field disrupts negative geotaxis in Drosophila via a CRY-dependent pathway. Nat Commun 5, 4391. doi: 10.1038/ncomms5391

34. Foley, L.E., Ling, J., Joshi, R., Evantal, N., Kadener, S., Emery, P., 2019. Drosophila PSI controls circadian period and the phase of circadian behavior under temperature cycle via tim splicing. eLife 8: e50063, doi: 10.7554/eLife.50063

35. Giesecke, A., Johnstone, P.S., Lamaze, A., Landskron, J., Atay, E., Chen, K.F., Wolf, E., Top, D., Stanewsky, R., 2023. A novel period mutation implicating nuclear export in temperature compensation of the Drosophila circadian clock. Curr Biol 33, 336–350, e5. doi: 10.1016/j.cub.2022.12.011

36. Gilbert, C., Maumus, F., 2022. Multiple Horizontal Acquisitions of Plant Genes in the Whitefly Bemisia tabaci. Genome Biol Evol 14, evac141. doi: 10.1093/gbe/evac141

37. Glossop, N.R., Lyons, L.C., Hardin, P.E., 1999. Interlocked feedback loops within the Drosophila circadian oscillator. Science 286, 766–768. doi: 10.1126/science.286.5440.766

38. Godinho, S.I., Maywood, E.S., Shaw, L., Tucci, V., Barnard, A.R., Busino, L., Pagano, M., Kendall, R., Quwailid, M.M., Romero, M.R., O’Neill, J., Chesham, J.E., Brooker, D., Lalanne, Z., Hastings, M.H., Nolan, P.M., 2007. The after-hours mutant reveals a role for Fbxl3 in determining mammalian circadian period. Science 316, 897–900. doi: 10.1126/science.1141138

39. Grima, B., Lamouroux, A., Chelot, E., Papin, C., Limbourg-Bouchon, B., Rouyer, F., 2002. The F-box protein slimb controls the levels of clock proteins period and timeless. Nature 420, 178–182. doi: 10.1038/nature01122

40. Hall, J.C., 2003. Genetics and molecular biology of rhythms in Drosophila and other insects. Adv Genet 48, 1–280. doi: 10.1016/s0065-2660(03)48000-0

41. Hardin, P.E., 2011. Molecular Genetic Analysis of Circadian Timekeeping in Drosophila. Adv Genet 74, 141–173. doi: 10.1016/B978-0-12-387690-4.00005-2

42. Hejnikova, M., Nouzova, M., Ramirez, C.E., Fernandez-Lima, F., Noriega, F.G., Dolezel, D., 2022. Sexual dimorphism of diapause regulation in the hemipteran bug Pyrrhocoris apterus. Insect Biochem Mol Biol 142, 103721. doi: 10.1016/j.ibmb.2022.103721

43. Helfrich-Förster, C., 2019. Light input pathways to the circadian clock of insects with an emphasis on the fruit fly Drosophila melanogaster. J Comp Physiol A Neuroethol Sens Neural Behav Physiol 206, 259–272. doi: 10.1007/s00359-019-01379-5

44. Helfrich-Forster, C., Winter, C., Hofbauer, A., Hall, J.C., Stanewsky, R., 2001. The circadian clock of fruit flies is blind after elimination of all known photoreceptors. Neuron 30, 249–261. doi: 10.1016/s0896-6273(01)00277-x

45. Hirano, A., Yumimoto, K., Tsunematsu, R., Matsumoto, M., Oyama, M., Kozuka-Hata, H., Nakagawa, T., Lanjakornsiripan, D., Nakayama, K.I., Fukada, Y., 2013. FBXL21 Regulates Oscillation of the Circadian Clock through Ubiquitination and Stabilization of Cryptochromes. Cell 152, 1106–1118. doi: 10.1016/j.cell.2013.01.054

46. Iiams, S.E., Wan, G., Zhang, J., Lugena, A.B., Zhang, Y., Hayden, A.N., Merlin, C., 2024. Loss of functional cryptochrome 1 reduces robustness of 24-hour behavioral rhythms in monarch butterflies. iScience 27, 108980. doi: 10.1016/j.isci.2024.108980

47. Ikeno, T., Ishikawa, K., Numata, H., Goto, S.G., 2013. Circadian clock gene Clock is involved in the photoperiodic response of the bean bug Riptortus pedestris. Physiol Entomol 38, 157–162. doi: 10.1111/Phen.12013

48. Ikeno, T., Katagiri, C., Numata, H., Goto, S.G., 2011a. Causal involvement of mammalian-type cryptochrome in the circadian cuticle deposition rhythm in the bean bug Riptortus pedestris. Insect Mol Biol 20, 409–415. doi: 10.1111/j.1365-2583.2011.01075.x

49. Ikeno, T., Numata, H., Goto, S.G., 2008. Molecular characterization of the circadian clock genes in the bean bug, Riptortus pedestris, and their expression patterns under long– and short-day conditions. Gene 419, 56–61. doi: 10.1016/j.gene.2008.05.002

50. Ikeno, T., Numata, H., Goto, S.G., 2011b. Circadian clock genes period and cycle regulate photoperiodic diapause in the bean bug Riptortus pedestris males. J Insect Physiol 57, 935–938. doi: 10.1016/j.jinsphys.2011.04.006

51. Ikeno, T., Numata, H., Goto, S.G., 2011c. Photoperiodic response requires mammalian-type cryptochrome in the bean bug Riptortus pedestris. Biochem Biophys Res Commun 410, 394–397. doi: 10.1016/j.bbrc.2011.05.142

52. Ikeno, T., Tanaka, S.I., Numata, H., Goto, S.G., 2010. Photoperiodic diapause under the control of circadian clock genes in an insect. BMC Biol 8, 116. doi: 10.1186/1741-7007-8-116

53. Jaumouille, E., Machado Almeida, P., Stahli, P., Koch, R., Nagoshi, E., 2015. Transcriptional Regulation via Nuclear Receptor Crosstalk Required for the Drosophila Circadian Clock. Curr Biol 25, 1502–1508. doi: 10.1016/j.cub.2015.04.017

54. Johnson, K.P., Dietrich, C.H., Friedrich, F., Beutel, R.G., Wipfler, B., Peters, R.S., Allen, J.M., Petersen, M., Donath, A., Walden, K.K.O., Kozlov, A.M., Podsiadlowski, L., Mayer, C., Meusemann, K., Vasilikopoulos, A., Waterhouse, R.M., Cameron, S.L., Weirauch, C., Swanson, D.R., Percy, D.M., Hardy, N.B., Terry, I., Liu, S., Zhou, X., Misof, B., Robertson, H.M., Yoshizawa, K., 2018. Phylogenomics and the evolution of hemipteroid insects. Proc Natl Acad Sci U S A 115, 12775–12780. doi: 10.1073/pnas.1815820115

55. Joshi, R., Cai, Y.D., Xia, Y.L., Chiu, J.C., Emery, P., 2022. PERIOD Phosphoclusters Control Temperature Compensation of the Drosophila Circadian Clock. Front Physiol 13: 888262. doi: 10.3389/fphys.2022.888262

56. Kadener, S., Stoleru, D., McDonald, M., Nawathean, P., Rosbash, M., 2007. Clockwork Orange is a transcriptional repressor and a new Drosophila circadian pacemaker component. Gene Dev 21, 1675–1686. doi: 10.1101/gad.1552607

57. Kamae, Y., Tomioka, K., 2012. timeless is an essential component of the circadian clock in a primitive insect, the firebrat Thermobia domestica. J Biol Rhythms 27, 126–134. doi: 10.1177/0748730411435997

58. Kamae, Y., Uryu, O., Miki, T., Tomioka, K., 2014. The Nuclear Receptor Genes HR3 and E75 Are Required for the Circadian Rhythm in a Primitive Insect. PLoS One 9, e114899. doi: 10.1371/journal.pone.0114899

59. Kaniewska, M.M., Chvalova, D., Dolezel, D., 2024. Impact of photoperiod and functional clock on male diapause in cryptochrome and pdf mutants in the linden bug Pyrrhocoris apterus. J Comp Physiol A Neuroethol Sens Neural Behav Physiol 210, 575–584. doi: 10.1007/s00359-023-01647-5

60. Kaniewska, M.M., Vaneckova, H., Dolezel, D., Kotwica-Rolinska, J., 2020. Light and Temperature Synchronizes Locomotor Activity in the Linden Bug, Pyrrhocoris apterus. Front Physiol 11, 242. doi: 10.3389/fphys.2020.00242

61. Kivimäe, S., Saez, L., Young, M.W., 2008. Activating PER repressor through a DBT-directed phosphorylation switch. PLoS Biol 6, 1570–1583. doi: 10.1371/journal.pbio.0060183

62. Ko, H.W., Jiang, J., Edery, I., 2002. Role for Slimb in the degradation of Drosophila Period protein phosphorylated by Doubletime. Nature 420, 673–678. doi: 10.1038/nature01272

63. Kobelkova, A., Bajgar, A., Dolezel, D., 2010. Functional Molecular Analysis of a Circadian Clock Gene timeless Promoter from the Drosophilid Fly Chymomyza costata. J Biol Rhythms 25, 399–409. doi: 10.1177/0748730410385283

64. Koh, K., Zheng, X., Sehgal, A., 2006. JETLAG resets the Drosophila circadian clock by promoting light-induced degradation of TIMELESS. Science 312, 1809–1812. doi: 10.1126/science.1124951

65. Komada, S., Kamae, Y., Koyanagi, M., Tatewaki, K., Hassaneen, E., Saifullah, A., Yoshii, T., Terakita, A., Tomioka, K., 2015. Green-sensitive opsin is the photoreceptor for photic entrainment of an insect circadian clock. Zoological Lett 1, 11. doi: 10.1186/s40851-015-0011-6

66. Kostal, V., 2011. Insect photoperiodic calendar and circadian clock: Independence, cooperation, or unity? J Insect Physiol 57, 538–556. doi: 10.1016/j.jinsphys.2010.10.006

67. Kotwica-Rolinska, J., Chodakova, L., Chvalova, D., Kristofova, L., Fenclova, I., Provaznik, J., Bertolutti, M., Wu, B.C., Dolezel, D., 2019. CRISPR/Cas9 Genome Editing Introduction and Optimization in the Non-model Insect Pyrrhocoris apterus. Front Physiol 10, 891. doi: 10.3389/fphys.2019.00891

68. Kotwica-Rolinska, J., Chodakova, L., Smykal, V., Damulewicz, M., Provaznik, J., Wu, B.C., Hejnikova, M., Chvalova, D., Dolezel, D., 2022a. Loss of Timeless Underlies an Evolutionary Transition within the Circadian Clock. Mol Biol Evol 39, msab346. doi: 10.1093/molbev/msab346

69. Kotwica-Rolinska, J., Damulewicz, M., Chodakova, L., Kristofova, L., Dolezel, D., 2022b. Pigment Dispersing Factor Is a Circadian Clock Output and Regulates Photoperiodic Response in the Linden Bug, Pyrrhocoris apterus. Front Physiol 13, 884909. doi: 10.3389/fphys.2022.884909

70. Kotwica-Rolinska, J., Pivarciova, L., Vaneckova, H., Dolezel, D., 2017. The role of circadian clock genes in the photoperiodic timer of the linden bug Pyrrhocoris apterus during the nymphal stage. Physiol Entomol 42, 266–273. doi: 10.1111/phen.12197

71. Kumar, M., Michael, S., Alvarado-Valverde, J., Meszaros, B., Samano-Sanchez, H., Zeke, A., Dobson, L., Lazar, T., Ord, M., Nagpal, A., Farahi, N., Kaser, M., Kraleti, R., Davey, N.E., Pancsa, R., Chemes, L.B., Gibson, T.J., 2022. The Eukaryotic Linear Motif resource: 2022 release. Nucleic Acids Res 50, D497–D508. doi: 10.1093/nar/gkab975

72. Kume, K., Zylka, M.J., Sriram, S., Shearman, L.P., Weaver, D.R., Jin, X.W., Maywood, E.S., Hastings, M.H., Reppert, S.M., 1999. mCRY1 and mCRY2 are essential components of the negative limb of the circadian clock feedback loop. Cell 98, 193–205. doi: 10.1016/s0092-8674(00)81014-4

73. Kutaragi, Y., Tokuoka, A., Tomiyama, Y., Nose, M., Watanabe, T., Bando, T., Moriyama, Y., Tomioka, K., 2018. A novel photic entrainment mechanism for the circadian clock in an insect: involvement of c-fos and cryptochromes. Zoological Lett 4, 26. doi: 10.1186/s40851-018-0109-8

74. Landskron, J., Chen, K.F., Wolf, E., Stanewsky, R., 2009. A Role for the PERIOD:PERIOD Homodimer in the Drosophila Circadian Clock. PLoS Biol 7, e1000003, 820–835. doi: 10.1371/journal.pbio.1000003

75. Lankinen, P., 1986. Geographical variation in circadian eclosion rhythm and photoperiodic adult diapause in Drosophila littoralis. J Comp Physiol 159:123–142. doi: 10.1007/BF00612503

76. Lankinen, P., 1993. North-south differences in circadian eclosion rhythm in European populations of Drosophila subobscura. Heredity 71, 210–218. doi: 10.1038/hdy.1993.126

77. Lapadula, W.J., Mascotti, M.L., Juri Ayub, M., 2020. Whitefly genomes contain ribotoxin coding genes acquired from plants. Sci Rep 10, 15503. doi: 10.1038/s41598-020-72267-1

78. Le Trionnaire, G., Tanguy, S., Hudaverdian, S., Gleonnec, F., Richard, G., Cayrol, B., Monsion, B., Pichon, E., Deshoux, M., Webster, C., Uzest, M., Herpin, A., Tagu, D., 2019. An integrated protocol for targeted mutagenesis with CRISPR-Cas9 system in the pea aphid. Insect Biochem Mol Biol 110, 34–44. doi: 10.1016/j.ibmb.2019.04.016

79. Lees, A.D., 1964. The Location of the Photoperiodic Receptors in the Aphid Megoura Viciae Buckton. J Exp Biol 41, 119–133. doi: 10.1242/jeb.41.1.119

80. Li, Y., Ma, S., Sun, L., Zhang, T., Chang, J., Lu, W., Chen, X., Liu, Y., Wang, X., Shi, R., Zhao, P., Xia, Q., 2018. Programmable Single and Multiplex Base-Editing in Bombyx mori Using RNA-Guided Cytidine Deaminases. G3 8, 1701–1709. doi: 10.1534/g3.118.200134

81. Ling, J., Dubruille, R., Emery, P., 2012. KAYAK-alpha Modulates Circadian Transcriptional Feedback Loops in Drosophila Pacemaker Neurons. J Neurosci 32, 16959–16970. doi: 10.1523/JNEUROSCI.1888-12.2012c

82. Majercak, J., Chen, W.F., Edery, I., 2004. Splicing of the period gene 3 ‘-terminal intron is regulated by light, circadian clock factors, and phospholipase C. Mol Cell Biol 24, 3359–3372. doi: 10.1128/MCB.24.8.3359-3372.2004

83. Majercak, J., Sidote, D., Hardin, P.E., Edery, I., 1999. How a circadian clock adapts to seasonal decreases in temperature and day length. Neuron 24: e44642, 219–230. doi: 10.1016/s0896-6273(00)80834-x

84. Martin Anduaga, A., Evantal, N., Patop, I.L., Bartok, O., Weiss, R., Kadener, S., 2019. Thermosensitive alternative splicing senses and mediates temperature adaptation in Drosophila. eLife 8. doi: 10.7554/eLife.44642

85. Meng, H., Gonzales, N.M., Jung, S.Y., Lu, Y., Putluri, N., Zhu, B., Dacso, C.C., Lonard, D.M., O’Malley, B.W., 2022. Defining the mammalian coactivation of hepatic 12-h clock and lipid metabolism. Cell Rep 38, 110491. doi: 10.1016/j.celrep.2022.110491

86. Merlin, C., Iiams, S.E., Lugena, A.B., 2020. Monarch Butterfly Migration Moving into the Genetic Era. Trends Genet 36, 689–701. doi: 10.1016/j.tig.2020.06.011

87. Mistry, J., Chuguransky, S., Williams, L., Qureshi, M., Salazar, G.A., Sonnhammer, E.L.L., Tosatto, S.C.E., Paladin, L., Raj, S., Richardson, L.J., Finn, R.D., Bateman, A., 2021. Pfam: The protein families database in 2021. Nucleic Acids Res 49, D412–D419. doi: 10.1093/nar/gkaa913

88. Mocchetti, A., De Rouck, S., Naessens, S., Dermauw, W., Van Leeuwen, T., 2024. SYNCAS based CRISPR-Cas9 gene editing in predatory mites, whiteflies and stinkbugs. Insect Biochem Mol Biol 177, 104232. doi: 10.1016/j.ibmb.2024.104232

89. Netusil, R., Tomanova, K., Chodakova, L., Chvalova, D., Dolezel, D., Ritz, T., Vacha, M., 2021. Cryptochrome-dependent magnetoreception in a heteropteran insect continues even after 24 h in darkness. J Exp Biol 224, jeb243000. doi: 10.1242/jeb.243000

90. Nguyen Ba, A.N., Pogoutse, A., Provart, N., Moses, A.M., 2009. NLStradamus: a simple Hidden Markov Model for nuclear localization signal prediction. BMC Bioinformatics 10, 202. doi: 10.1186/1471-2105-10-202

91. Ogueta, M., Hardie, R.C., Stanewsky, R., 2018. Non-canonical Phototransduction Mediates Synchronization of the Drosophila melanogaster Circadian Clock and Retinal Light Responses. Curr Biol 28, 1725–1735.e3 doi: 10.1016/j.cub.2018.04.016

92. Ozkaya, O., Rosato, E., 2012. The circadian clock of the fly: a neurogenetics journey through time. Adv Genet 77, 79–123. doi: 10.1016/B978-0-12-387687-4.00004-0

93. Ozturk, N., Vanvickle-Chavez, S.J., Akileswaran, L., Van Gelder, R.N., Sancar, A., 2013. Ramshackle (Brwd3) promotes light-induced ubiquitylation of Drosophila Cryptochrome by DDB1-CUL4-ROC1 E3 ligase complex. Proc Natl Acad Sci U S A 110, 4980–4985. doi: 10.1073/pnas.1303234110

94. Pacheco, I.D., Walling, L.L., Atkinson, P.W., 2022. Gene Editing and Genetic Control of Hemipteran Pests: Progress, Challenges and Perspectives. Front Bioeng Biotechnol 10, 900785. doi: 10.3389/fbioe.2022.900785

95. Paolucci, S., Dalla Benetta, E., Salis, L., Dolezel, D., van de Zande, L., Beukeboom, L.W., 2019. Latitudinal Variation in Circadian Rhythmicity in Nasonia vitripennis. Behav Sci 9. doi: 10.3390/bs9110115

96. Pavelka, J., Shimada, K., Kostal, V., 2003. TIMELESS: A link between fly’s circadian and photoperiodic clocks? Eur J Entomol 100, 255–265. doi: 10.14411/eje.2003.041

97. Peschel, N., Chen, K.F., Szabo, G., Stanewsky, R., 2009. Light-Dependent Interactions between the Drosophila Circadian Clock Factors Cryptochrome, Jetlag, and Timeless. Curr Biol 19, 241–247. doi: 10.1016/j.cub.2008.12.042

98. Pivarciova, L., Vaneckova, H., Provaznik, J., Wu, B.C., Pivarci, M., Peckova, O., Bazalova, O., Cada, S., Kment, P., Kotwica-Rolinska, J., Dolezel, D., 2016. Unexpected Geographic Variability of the Free Running Period in the Linden Bug Pyrrhocoris apterus. J Biol Rhythms 31, 568–576. doi: 10.1177/0748730416671213

99. Putker, M., Wong, D.C.S., Seinkmane, E., Rzechorzek, N.M., Zeng, A., Hoyle, N.P., Chesham, J.E., Edwards, M.D., Feeney, K.A., Fischer, R., Peschel, N., Chen, K.F., Vanden Oever, M., Edgar, R.S., Selby, C.P., Sancar, A., O’Neill, J.S., 2021. CRYPTOCHROMES confer robustness, not rhythmicity, to circadian timekeeping. EMBO J 40, e106745. doi: 10.15252/embj.2020106745

100. Reding, K., Pick, L., 2020. High-Efficiency CRISPR/Cas9 Mutagenesis of the white Gene in the Milkweed Bug Oncopeltus fasciatus. Genetics 215, 1027–1037. doi: 10.1534/genetics.120.303269

101. Richier, B., Michard-Vanhee, C., Lamouroux, A., Papin, C., Rouyer, F., 2008. The clockwork orange Drosophila protein functions as both an activator and a repressor of clock gene expression. J Biol Rhythms 23, 103–116. doi: 10.1177/0748730407313817

102. Rivas, G.B.S., Zhou, J., Merlin, C., Hardin, P.E., 2021. CLOCKWORK ORANGE promotes CLOCK-CYCLE activation via the putative Drosophila ortholog of CLOCK INTERACTING PROTEIN CIRCADIAN. Curr Biol 31:e4204, 4207–4218. doi: 10.1016/j.cub.2021.07.017

103. Rubin, E.B., Shemesh, Y., Cohen, M., Elgavish, S., Robertson, H.M., Bloch, G., 2006. Molecular and phylogenetic analyses reveal mammalian-like clockwork in the honey bee (Apis mellifera) and shed new light on the molecular evolution of the circadian clock. Genome Res 16, 1352–1365. doi: 10.1101/gr.5094806

104. Rutila, J.E., Suri, V., Le, M., So, W.V., Rosbash, M., Hall, J.C., 1998. CYCLE is a second bHLH-PAS clock protein essential for circadian rhythmicity and transcription of Drosophila period and timeless. Cell 93, 805–814. doi: 10.1016/s0092-8674(00)81441-5

105. Saez, L., Meyer, P., Young, M.W., 2007. A PER/TIM/DBT interval timer for Drosophila’s circadian clock. Cold Spring Harb Symp Quant Biol 72, 69–74. doi: 10.1101/sqb.2007.72.034

106. Saez, L., Young, M.W., 1996. Regulation of nuclear entry of the Drosophila clock proteins period and timeless. Neuron 17, 911–920. doi: 10.1016/s0896-6273(00)80222-6

107. Saint-Charles, A., Michard-Vanhee, C., Alejevski, F., Chelot, E., Boivin, A., Rouyer, F., 2016. Four of the six Drosophila rhodopsin-expressing photoreceptors can mediate circadian entrainment in low light. J Comp Neurol 524, 2828–2844. doi: 10.1002/cne.23994

108. Saunders, D.S., 2005. Erwin Bunning and Tony Lees, two giants of chronobiology, and the problem of time measurement in insect photoperiodism. J Insect Physiol 51, 599–608. doi: 10.1016/j.jinsphys.2004.12.002

109. Saunders, D.S., 2010. Controversial aspects of photoperiodism in insects and mites. J Insect Physiol 56, 1491–1502. doi: 10.1016/j.jinsphys.2010.05.002

110. Sehgal, A., Price, J.L., Man, B., Young, M.W., 1994. Loss of circadian behavioral rhythms and per RNA oscillations in the Drosophila mutant timeless. Science 263, 1603–1606. doi: 10.1126/science.8128246

111. Senthilan, P.R., Helfrich-Förster, C., 2016. Rhodopsin 7-The unusual Rhodopsin in Drosophila. PeerJ 4, e2427. doi: 10.7717/peerj.2427

112. Shakhmantsir, I., Nayak, S., Grant, G.R., Sehgal, A., 2018. Spliceosome factors target timeless (tim) mRNA to control clock protein accumulation and circadian behavior in Drosophila. eLife 7: e39821. doi: 10.7554/eLife.39821

113. Shirai, Y., Piulachs, M.D., Belles, X., Daimon, T., 2022. DIPA-CRISPR is a simple and accessible method for insect gene editing. Cell Rep Methods 2, 100215. doi: 10.1016/j.crmeth.2022.100215

114. Siepka, S.M., Yoo, S.H., Park, J., Song, W.M., Kumar, V., Hu, Y.N., Lee, C., Takahashi, J.S., 2007. Circadian mutant overtime reveals F-box protein FBXL3 regulation of cryptochrome and period gene expression. Cell 129, 1011–1023. doi: 10.1016/j.cell.2007.04.030

115. Singh, S., Giesecke, A., Damulewicz, M., Fexova, S., Mazzotta, G.M., Stanewsky, R., Dolezel, D., 2019. New Drosophila Circadian Clock Mutants Affecting Temperature Compensation Induced by Targeted Mutagenesis of Timeless. Front Physiol 10, 1442. doi: 10.3389/fphys.2019.01442

116. Smykal, V., Bajgar, A., Provaznik, J., Fexova, S., Buricova, M., Takaki, K., Hodkova, M., Jindra, M., Dolezel, D., 2014a. Juvenile hormone signaling during reproduction and development of the linden bug, Pyrrhocoris apterus. Insect Biochem Mol Biol 45, 69–76. doi: 10.1016/j.ibmb.2013.12.003

117. Smykal, V., Chodakova, L., Hejnikova, M., Briedikova, K., Wu, B.C., Vaneckova, H., Chen, P., Janovska, A., Kyjakova, P., Vacha, M., Dolezel, D., 2023. Steroid receptor coactivator TAIMAN is a new modulator of insect circadian clock. PLoS Genet 19, e1010924. doi: 10.1371/journal.pgen.1010924

118. Smykal, V., Daimon, T., Kayukawa, T., Takaki, K., Shinoda, T., Jindra, M., 2014b. Importance of juvenile hormone signaling arises with competence of insect larvae to metamorphose. Dev Biol 390, 221–230. doi: 10.1016/j.ydbio.2014.03.006

119. Smykal, V., Dolezel, D., 2023. Evolution of proteins involved in the final steps of juvenile hormone synthesis. J Insect Physiol 145, 104487. doi: 10.1016/j.jinsphys.2023.104487

120. Smykal, V., Pivarci, M., Provaznik, J., Bazalova, O., Jedlicka, P., Luksan, O., Horak, A., Vaneckova, H., Benes, V., Fiala, I., Hanus, R., Dolezel, D., 2020. Complex Evolution of Insect Insulin Receptors and Homologous Decoy Receptors, and Functional Significance of Their Multiplicity. Mol Biol Evol 37, 1775–1789. doi: 10.1093/molbev/msaa048

121. Stashi, E., Lanz, R.B., Mao, J., Michailidis, G., Zhu, B., Kettner, N.M., Putluri, N., Reineke, E.L., Reineke, L.C., Dasgupta, S., Dean, A., Stevenson, C.R., Sivasubramanian, N., Sreekumar, A., Demayo, F., York, B., Fu, L., O’Malley, B.W., 2014. SRC-2 is an essential coactivator for orchestrating metabolism and circadian rhythm. Cell Rep 6, 633–645. doi: 10.1016/j.celrep.2014.01.027

122. Stehlik, J., Zavodska, R., Shimada, K., Sauman, I., Kostal, V., 2008. Photoperiodic induction of diapause requires regulated transcription of timeless in the larval brain of Chymomyza costata. J Biol Rhythms 23, 129–139. doi: 10.1177/0748730407313364

123. Takeuchi, K., Matsuka, M., Shinohara, T., Hamada, M., Tomiyama, Y., Tomioka, K., 2023. Fbxl4 Regulates the Photic Entrainment of Circadian Locomotor Rhythms in the Cricket Gryllus bimaculatus. Zoolog Sci 40, 53–63. doi: 10.2108/zs220047

124. Thakkar, N., Giesecke, A., Bazalova, O., Martinek, J., Smykal, V., Stanewsky, R., Dolezel, D., 2022. Evolution of casein kinase 1 and functional analysis of new doubletime mutants in Drosophila. Front Physiol 13, 1062632. doi: 10.3389/fphys.2022.1062632

125. Thakkar, N., Hejzlarova, A., Brabec, V., Dolezel, D., 2023. Germline Editing of Drosophila Using CRISPR-Cas9-based Cytosine and Adenine Base Editors. CRISPR J 6: 557–569. doi: 10.1089/crispr.2023.0026

126. Tobita, H., Kiuchi, T., 2024. Knockout of cryptochrome 1 disrupts circadian rhythm and photoperiodic diapause induction in the silkworm, Bombyx mori. Insect Biochem Mol Biol 172, 104153. doi: 10.1016/j.ibmb.2024.104153

127. Tomioka, K., 2014. Chronobiology of crickets: a review. Zoolog Sci 31, 624–632. doi: 10.2108/zs140024

128. Tomioka, K., Matsumoto, A., 2015. Circadian molecular clockworks in non-model insects. Curr Opin Insect Sci 7, 58–64. doi: 10.1016/j.cois.2014.12.006

129. Tomioka, K., Matsumoto, A., 2019. The circadian system in insects: Cellular, molecular, and functional organization. Advances in Insect Physiology, Vol 56, 73–115. doi: 10.1016/bs.aiip.2019.01.001

130. Tumova, S., Dolezel, D., Jindra, M., 2024. Conserved and Unique Roles of bHLH-PAS Transcription Factors in Insects – From Clock to Hormone Reception. J Mol Biol 436 (2024) 168332, 1-25. doi: 10.1016/j.jmb.2023.168332

131. Urbanova, V., Bazalova, O., Vaneckova, H., Dolezel, D., 2016. Photoperiod regulates growth of male accessory glands through juvenile hormone signaling in the linden bug, Pyrrhocoris apterus. Insect Biochem Mol Biol 70, 184–190. doi: 10.1016/j.ibmb.2016.01.003

132. Vafopoulou, X., Steel, C.G., 2012. Metamorphosis of a clock: remodeling of the circadian timing system in the brain of Rhodnius prolixus (Hemiptera) during larval-adult development. J Comp Neurol 520, 1146–1164. doi: 10.1002/cne.22743

133. Vafopoulou, X., Steel, C.G., Terry, K.L., 2007. Neuroanatomical relations of prothoracicotropic hormone neurons with the circadian timekeeping system in the brain of larval and adult Rhodnius prolixus (Hemiptera). J Comp Neurol 503, 511–524. doi: 10.1002/cne.21393

134. Wan, G., Hayden, A.N., Iiams, S.E., Merlin, C., 2021. Cryptochrome 1 mediates light-dependent inclination magnetosensing in monarch butterflies. Nat Commun 12, 771. doi: 10.1038/s41467-021-21002-z

135. Wan, G., Liu, R., Li, C., He, J., Pan, W., Sword, G.A., Hu, G., Chen, F., 2020. Change in geomagnetic field intensity alters migration-associated traits in a migratory insect. Biol Lett 16, 20190940. doi: 10.1098/rsbl.2019.0940

136. Werckenthin, A., Huber, J., Arnold, T., Koziarek, S., Plath, M.J.A., Plath, J.A., Stursberg, O., Herzel, H., Stengl, M., 2020. Neither per, nor tim1, nor cry2 alone are essential components of the molecular circadian clockwork in the Madeira cockroach. PLoS One 15, e0235930. doi: 10.1371/journal.pone.0235930

137. Xi, J., Toyoda, I., Shiga, S., 2017. Afferent neural pathways from the photoperiodic receptor in the bean bug, Riptortus pedestris. Cell Tissue Res. 368: 469–485 doi: 10.1007/s00441-016-2565-9

138. Xia, J., Guo, Z., Yang, Z., Han, H., Wang, S., Xu, H., Yang, X., Yang, F., Wu, Q., Xie, W., Zhou, X., Dermauw, W., Turlings, T.C.J., Zhang, Y., 2021. Whitefly hijacks a plant detoxification gene that neutralizes plant toxins. Cell 184, 1693–1705. doi: 10.1016/j.cell.2021.02.014, Erratum in: Cell. 2021 184: 3588. doi: 10.1016/j.cell.2021.06.010

139. Xue, W.H., Xu, N., Yuan, X.B., Chen, H.H., Zhang, J.L., Fu, S.J., Zhang, C.X., Xu, H.J., 2018. CRISPR/Cas9-mediated knockout of two eye pigmentation genes in the brown planthopper, Nilaparvata lugens (Hemiptera: Delphacidae). Insect Biochem Mol Biol 93, 19–26. doi: 10.1016/j.ibmb.2017.12.003

140. Yoshii, T., Ahmad, M., Helfrich-Forster, C., 2009. Cryptochrome Mediates Light-Dependent Magnetosensitivity of Drosophila’s Circadian Clock. PLoS Biol 7, 813–819. doi: 10.1371/journal.pbio.1000086

141. Yuan, Q., Metterville, D., Briscoe, A.D., Reppert, S.M., 2007. Insect cryptochromes: Gene duplication and loss define diverse ways to construct insect circadian clocks. Mol Biol Evol 24, 948–955. doi: 10.1093/molbev/msm011

142. Zhang, Y., Markert, M.J., Groves, S.C., Hardin, P.E., Merlin, C., 2017. Vertebrate-like CRYPTOCHROME 2 from monarch regulates circadian transcription via independent repression of CLOCK and BMAL1 activity. Proc Natl Acad Sci U S A 114, E7516–E7525. doi: 10.1073/pnas.1702014114

143. Zhou, J., Yu, W., Hardin, P.E., 2016. CLOCKWORK ORANGE Enhances PERIOD Mediated Rhythms in Transcriptional Repression by Antagonizing E-box Binding by CLOCK-CYCLE. PLoS Genet 12: e1006430. doi: 10.1371/journal.pgen.1006430

