## Supplementary material for "Evolution of circadian clock and light-input pathway genes in Hemiptera": upplementary material:

Short title:
Hemipteriod circadian clock

Authors: Vlastimil Smykal[1,]*, Hisashi Tobita[1,2], and David Dolezel[1,]*

**Affiliations:**

[1]Biology Centre of the Czech Academy of Sciences; České Budějovice, 37005, Czech Republic

[2]present address: Department of Agricultural and Environmental Biology, Graduate School of Agricultural and Life Sciences, The University of Tokyo, Bunkyo-ku, Tokyo, 113-8657, Japan

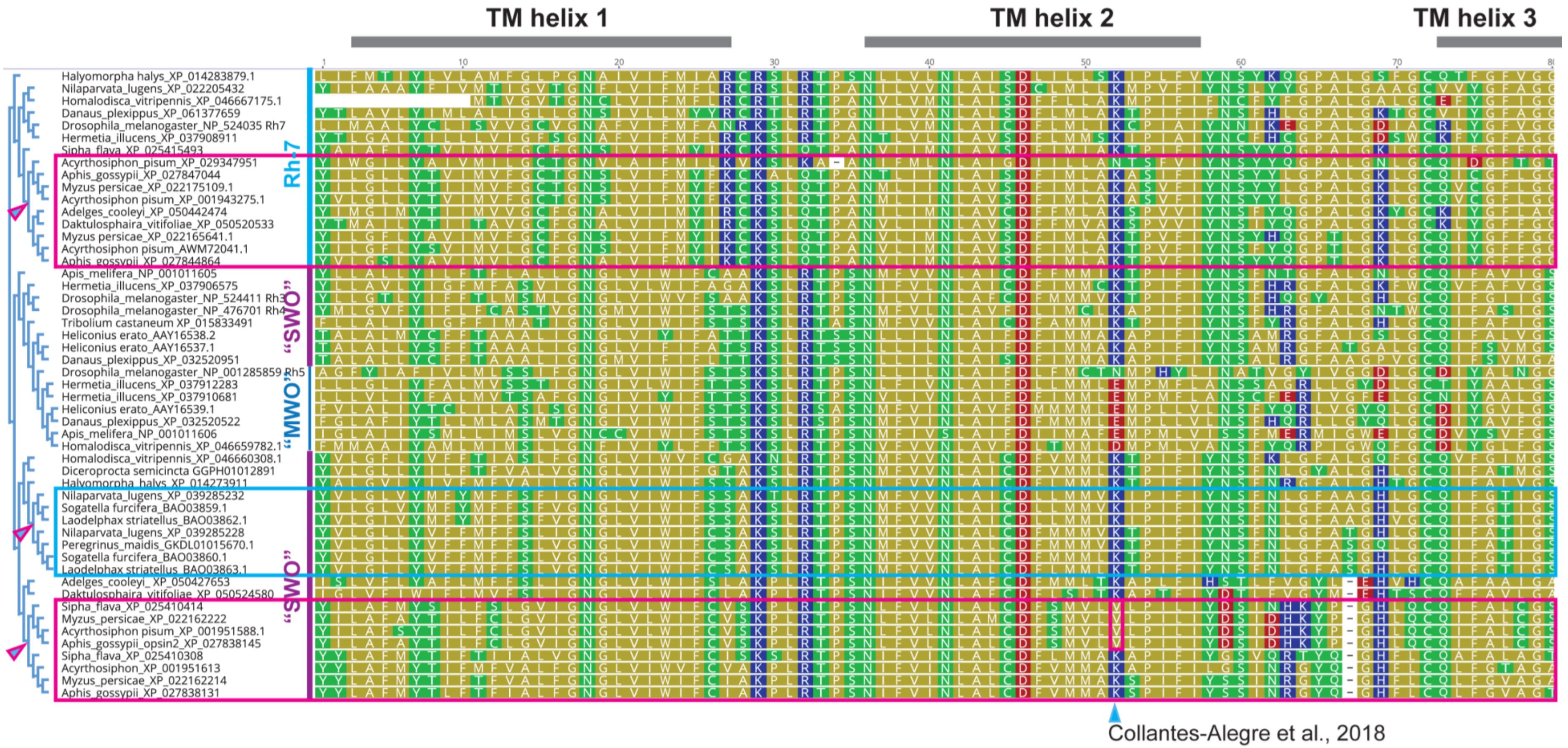


***opsin align page1 2025-01-15b***

**Fig. S1A.** Protein alignment of selected opsins from the Rh7, SWO, and MWO clades. The phylogenetic tree is shown on the far left, with arrowheads indicating gene duplication events. These duplications are marked with rectangles in the alignment. Aphid-specific duplications are highlighted in magenta, while the planthopper-specific duplication of SWO is highlighted in turquoise. TM helix refers to the transmembrane α-helix. The arrowhead at the bottom indicates a substitution in SWO that is suggested to potentially impact the spectral properties of opsins.

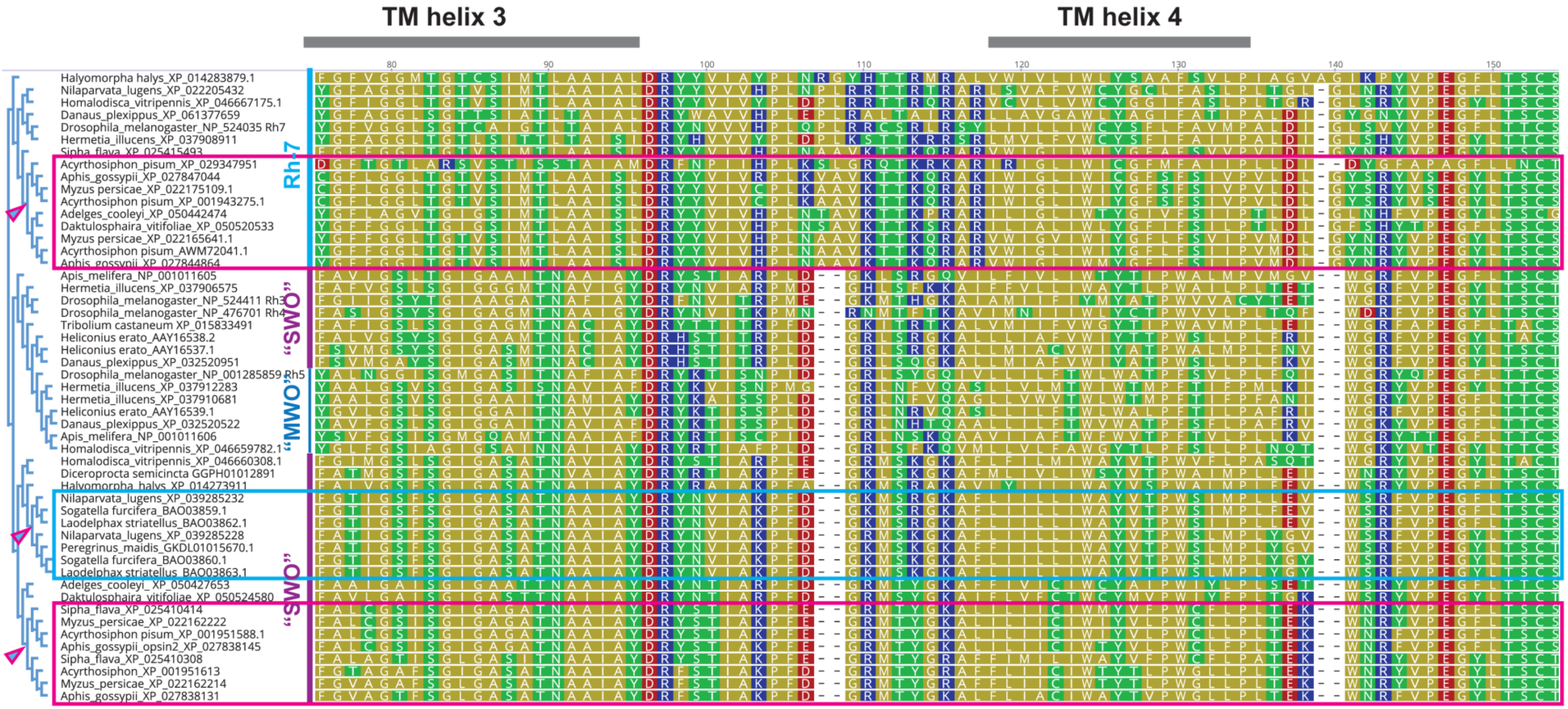


*opsin align page2 2025-01-15b*

**Fig. S1B.** Protein alignment of selected opsins from the Rh7, SWO, and MWO clades. The phylogenetic tree is shown on the far left, with arrowheads indicating gene duplication events. These duplications are marked with rectangles in the alignment. Aphid-specific duplications are highlighted in magenta, while the planthopper-specific duplication of SWO is highlighted in turquoise. TM helix refers to the transmembrane α-helix.

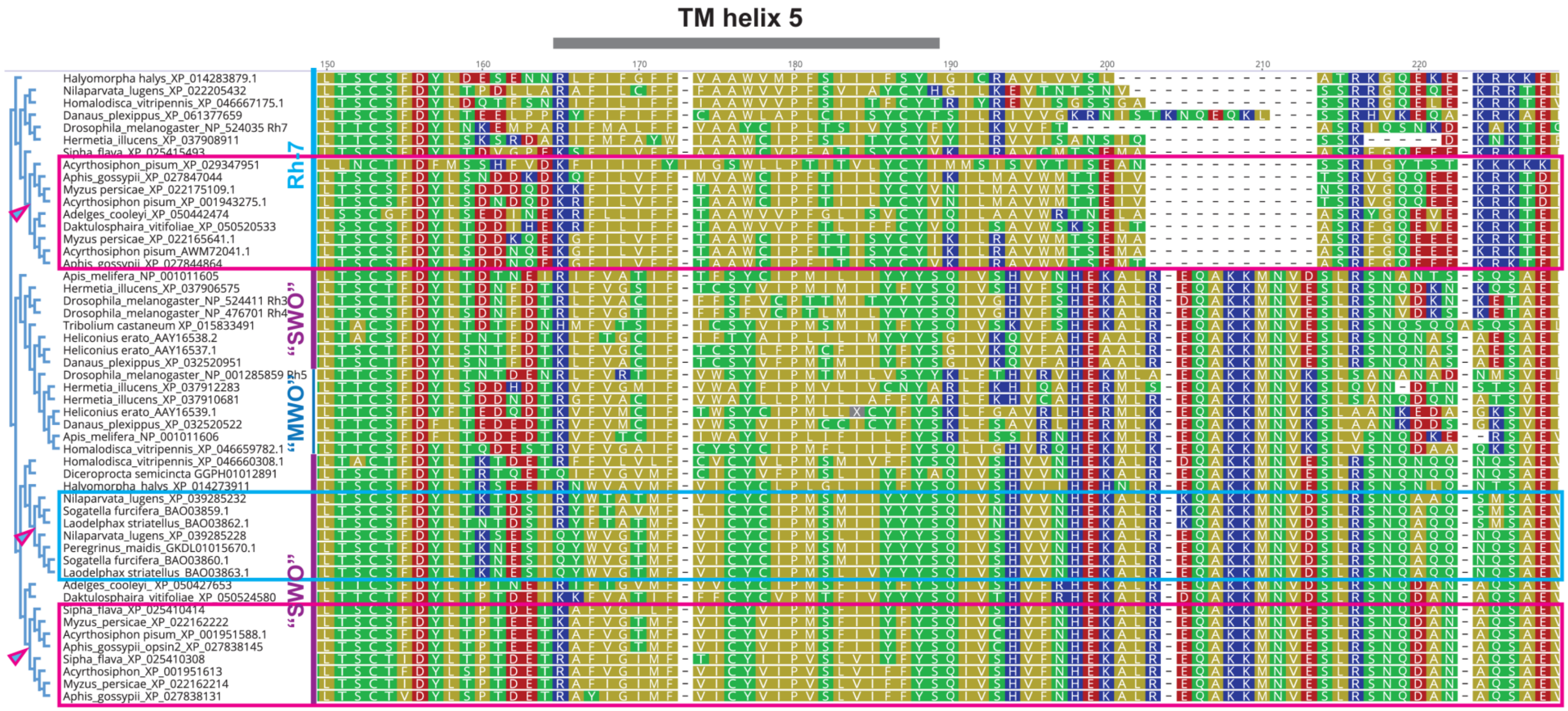


***opsin align page3 2025-01-15b***

**Fig. S1C.** Protein alignment of selected opsins from the Rh7, SWO, and MWO clades. The phylogenetic tree is shown on the far left, with arrowheads indicating gene duplication events. These duplications are marked with rectangles in the alignment. Aphid-specific duplications are highlighted in magenta, while the planthopper-specific duplication of SWO is highlighted in turquoise. TM helix refers to the transmembrane α-helix.

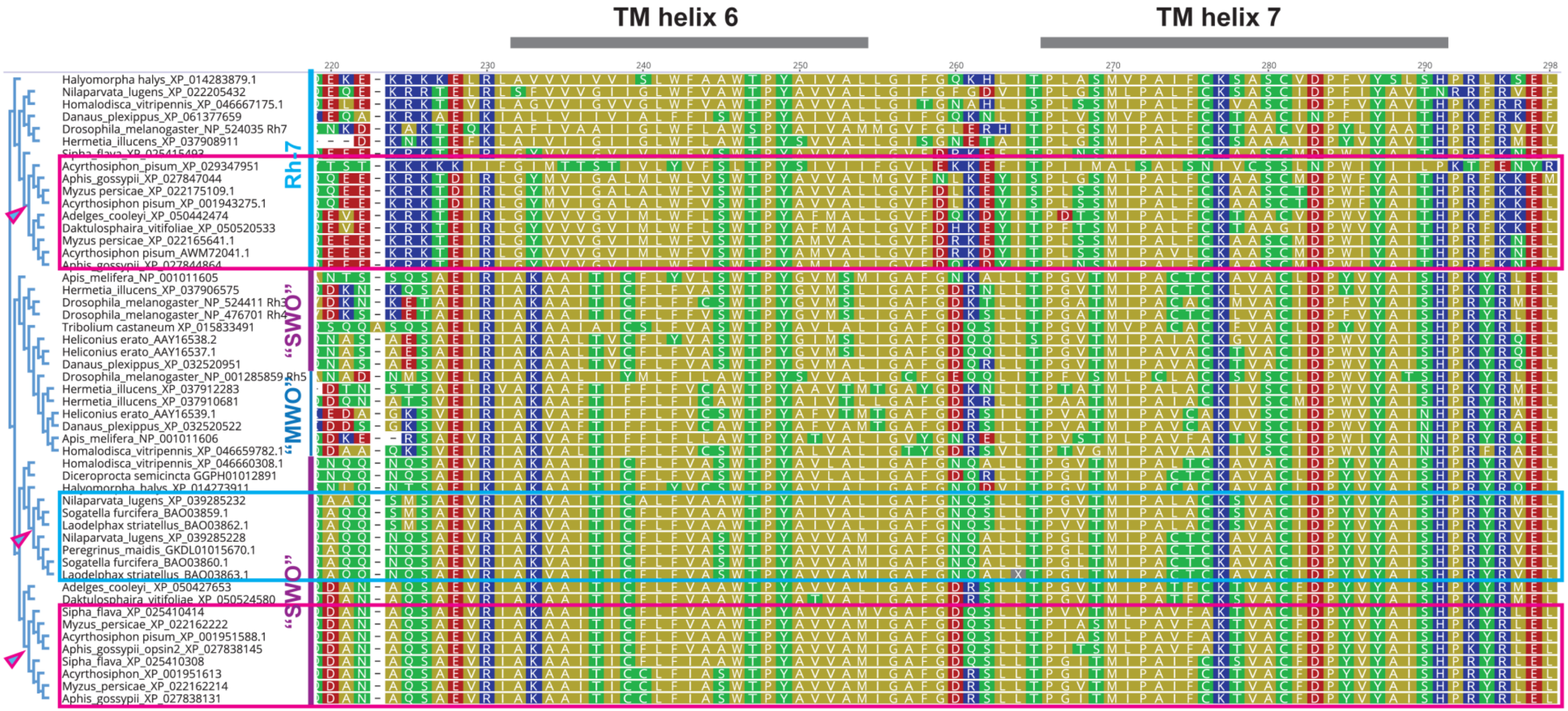


***opsin align page4 2025-01-15b***

**Fig. S1D.** Protein alignment of selected opsins from the Rh7, SWO, and MWO clades. The phylogenetic tree is shown on the far left, with arrowheads indicating gene duplication events. These duplications are marked with rectangles in the alignment. Aphid-specific duplications are highlighted in magenta, while the planthopper-specific duplication of SWO is highlighted in turquoise. TM helix refers to the transmembrane α-helix.

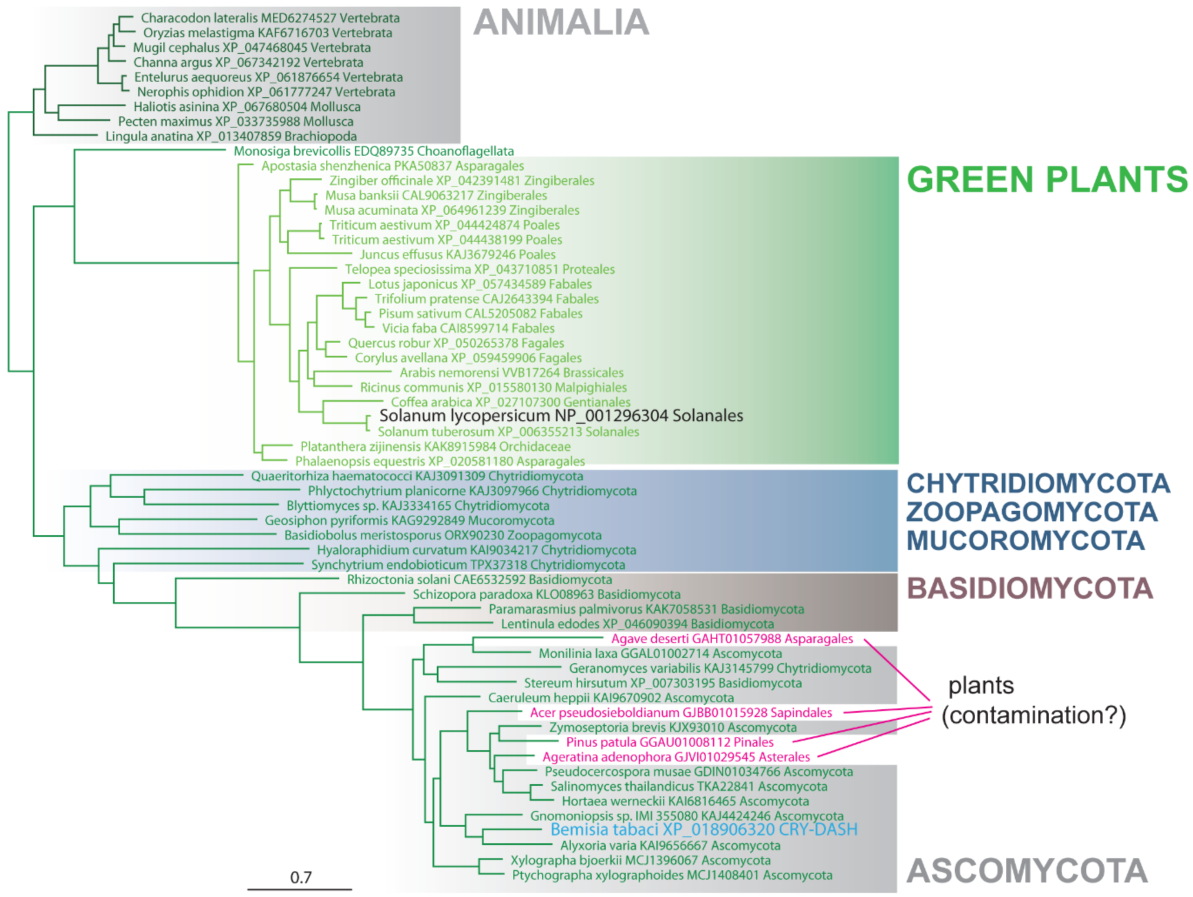


***SI DASHtree 2025-01-16b***

**Fig. S2** Phylogenetic analysis of CRY-DASH proteins. Animal CRY-DASH proteins served as an outgroup (top, Animalia). One cluster contains only CRY-DASH from green plants (green background, middle part). The bottom branch includes several representatives of fungal lineages (Chytridiomycota, Mucoromycota, Zoopagomycota, Basidiomycota, and Ascomycota), highlighted with different background colors. The *Bemisia* CRY-DASH branches within *Ascomycota*. Four plant "CRY-DASH" protein sequences branching within fungi were identified in the Transcriptome Shotgun Assembly (TSA) databases. These sequences are highly suspicious and are likely the result of cross-contamination of the plant sample with fungi.

**Table S1.** Synteny in *(6-4)-photolyase* locus

| species + contig acc # | Gene or cDNA/TSA, species, acc # |
|---|---|
| *Homalodisca vitripennis* | *(6-4)-photolyase* (*6-4 ph*) 124364814 gene |
| *Homalodisca vitripennis* NC_060212.1 | *Transducer of ERBB2* (*Btg3-like*) 124364807 gene |
| | *Cyclin-Y* (*CycY*) 124364810 gene |
| | *Golgin subfamily A member 2* (*Golga2*) 124364811 gene |
| | *Phosphotyrosyl phosphatase activator = Serine/threonine-protein phosphatase 2A activator* (*Ptpa*) 124364813 gene |
| | *Heparan sulfate 2-O-sulfotransferase* (*Hs2st*) 124364816 gene |
| | *Multiple epidermal growth factor-like domains protein 8* (*Megf8*) 124364817 gene |
| | *Venom dipeptidyl peptidase 4* (*Vdpp4*) 124364820 gene |
| | *Nucleolar protein 10* (*Nop10*) 124364822 gene |
| *Nilaparvata lugens* | |
| *Nilaparvata lugens* NC_052506.1 | *Btg3-like* GANM01011305.1 |
| | *CycY* GAYF02024452.1 |
| | *Golga2* GANM01000410.1 |
| | *Ptpa* GAYF02022189.1 |
| | *Hs2st* GAYF02023552.1 |
| | *Megf8* IACV01042805.1 |
| | *Vdpp4* IACV01064837.1 |
| | *Nop10* GAYF02024693.1 |
| *Laodelphax striatellus* | |
| *Laodelphax striatellus* JADWYY010000002.1 | *Btg3-like* 111044182 gene |
| | *CycY* 111050054 gene |
| | *Golga2* 111062024 gene |
| | *Ptpa* 111058596 gene |
| | *Hs2st* 111043809 gene |
| | *Megf8* 111057596 gene |
| | *Vdpp4* 111052236 gene |
| | *Nop10* 111055520 gene |

**Table S2.** *CRY-DASH* synteny

| species + contig acc # | Gene or cDNA/TSA, species, acc # |
|---|---|
| *Bemisia tabaci* | *DASH* (DASH) 109036507 gene |
| *Bemisia tabaci* NW_017548003.1 | *uridine 5'-monophosphate synthase* (*UPM synthase*) 109036513 gene |
| | *DNA repair protein complementing XP-A cells* (*xpa*) 109036503 gene |
| | *peroxisomal membrane protein 11B* (*Pex11B*) 109036502 gene |
| | *transient receptor potential-gamma protein* (*Trp gamma*) 109036537 gene |
| | *U8-agatoxin-Ao1a-like* (*TXAG8*) 109036525 gene |
| | *peptidyl-prolyl cis-trans isomerase, rhodopsin-specific isozyme-like* (*ninaA*) 109036508 gene |
| | *Zwilch* (*Zwilch*) 109036515 gene |
| *Trialeurodes vaporariorum* | |
| *Trialeurodes vaporariorum* VMOF01000024.1 | *UPM synthase* TSA: GHMB01019681.1 |
| | *xpa* TSA: GHMB01073600.1 |
| | *Pex11B* TSA: GHMB01078043.1 |
| | *Trp gamma* TSA: GHMB01034226.1 |
| | *TXAG8* TSA: GHMB01051084.1 |
| | *Zwilch* TSA: GHMB01009679.1 |
| *Trialeurodes vaporariorum* VMOF01000035.1 | ninaA TSA: GHMB01074035.1 |
| *Solanum lycopersicum* | |
| *Solanum lycopersicum JALGYV010003334.1* | |
